# Variation in the transcriptome response and detoxification gene diversity drives pesticide tolerance in fishes

**DOI:** 10.1101/2021.12.16.473024

**Authors:** M.J. Lawrence, P. Grayson, J.D. Jeffrey, M.F. Docker, C.J. Garroway, J.M. Wilson, R.G. Manzon, M.P. Wilkie, K.M. Jeffries

## Abstract

Pesticides are critical for invasive species management, but often have negative effects on non-target native biota. Tolerance to pesticides should have an evolutionary basis, but this is poorly understood. Invasive sea lamprey (*Petromyzon marinus*) populations in North America have been controlled with a pesticide lethal to them at lower concentrations than native fishes. We addressed how interspecific variation in gene expression and detoxification gene diversity confer differential pesticide sensitivity in two fish species. We exposed sea lamprey and bluegill (*Lepomis macrochirus*), a tolerant native species, to TFM, a pesticide commonly used in sea lamprey control, and used whole-transcriptome sequencing of gill and liver to characterize the cellular response. Comparatively, bluegill exhibited a larger number of detoxification genes expressed and a larger number of responsive transcripts overall, which likely contributes to greater tolerance to TFM. Understanding the genetic and physiological basis for pesticide tolerance is crucial for managing invasive species.

---

Invasive species represent a considerable threat to local biodiversity and ecosystem functioning, having significant economic costs associated with lost ecosystem services and control efforts^1^. Invasive species control is multifaceted and serves primarily to reduce, but not necessarily eradicate, populations of the target organism^2^. Often, chemical pesticides are used to kill the target species while attempting to minimize non-target impacts^2^. In aquatic environments, this can be difficult as pesticides are often applied to large portions of the waterbody, thus exposing all community members^3^. However, species-specific pesticides are relatively rare and, consequently, there will likely be impacts on native species in most pesticide application programs^4^.

The degree to which different species are adversely affected by pesticide applications can be highly species-specific. Interspecific variation in pesticide sensitivities occurs in amphibians^5^, invertebrates^6^, and fishes^7^, which suggests that species-specific differences in pesticide uptake, biotransformation, and elimination (i.e., the disposition) are likely driving differential sensitivity^8^. Further, phylogenetic signatures associated with toxicity tolerances have been identified in several clades of aquatic ectotherms^9^. This suggests that toxicity tolerances likely have an evolutionary basis and are in part genetically determined^10^. Recent transcriptional profiling of several teleost fishes has shown considerable interspecific variation amongst genes involved in detoxification of pesticides and other xenobiotics^11^, underscoring an important genomic role in toxicity tolerances. Consequently, using molecular toolsets for understanding pesticide responses are valuable in providing insight into phylogenetic variation in physiological response and in predicting community impacts of pesticide applications^9, 10^. This is relevant in contemporary invasion control efforts, where minimizing harm to native species is often a key objective^12^. To date, our understanding of the genetic and evolutionary underpinnings of interspecific variation in toxicity tolerance is limited, hindering our insight into community-level effects of pesticide applications.

The sea lamprey (*Petromyzon marinus*) control program in the Laurentian Great Lakes of North America (hereafter referred to as the Great Lakes) is an excellent model system to explore the evolutionary and genetic foundations of differential toxicity tolerance to a pesticide in target (i.e., invasive sea lamprey) and non-target (native) species. Sea lamprey, although native to the Atlantic Ocean, likely gained access to Lake Ontario through canals in the mid-1800s; the Welland Canal subsequently allowed access to the remaining Great Lakes by the early 1900s^13^. By the 1950s, sea lamprey populations in the Great Lakes had exploded, decimating native fisheries, notably lake trout (*Salvelinus namaycush*), having severe impacts on the environment and economy of the region^13^. An international Great Lakes Fishery Commission (GLFC) involving Canadian and American institutions was formed in 1955 to co-manage fisheries and to control sea lamprey populations. Since the 1950s, the GFLC’s primary method of sea lamprey control has involved treating streams and rivers with pesticides, commonly referred to as lampricides, that target burrowing sea lamprey larvae in Great Lakes tributaries^13^. Because the filter-feeding larval stage lasts for ∼4–5 years before sea lamprey metamorphose into parasitic juveniles that disperse widely in the lakes, lampricide treatments in streams containing sea lamprey larvae kill multiple year classes at once.

Two pesticides are used to control sea lamprey, 3-trifluoromethyl-4-nitrophenol (TFM) and niclosamide (2’,5-dichloro-4’-nitrosalicylanilide), each of which contains a halogenated phenol ring that contributes to their toxicity^3^. In most treatments, TFM is applied at concentrations reaching the sea lamprey 9-h LC99.9 of TFM^14^, the concentration needed to kill 99.9% of the exposed population, and is co-applied with 1-2% niclosamide because these compounds interact in a greater than additive fashion^15^. While niclosamide is broadly toxic to fishes^3^, TFM is highly toxic to lampreys relative to jawed fishes. The mode of action of TFM is interference with oxidative phosphorylation in the mitochondria^16^, which lowers aerobic ATP production forcing the animal to increasingly rely on unsustainable anaerobic glycolysis^17–19^. However, TFM sensitivity varies greatly among jawed fishes, which makes understanding the broader non-target impacts of TFM difficult to resolve^3^. For example, lake sturgeon (*Acipenser fulvescens*), a non-teleost jawed fish, are moderately sensitive to TFM^3^, while centrarchids, which includes basses and bluegill (*Lepomis macrochirus*), have some of the highest reported tolerances to TFM amongst teleost fishes in the Great Lakes, with reported LC values ∼10-fold higher than sea lamprey^19^.

Little is understood about the physiological mechanisms driving varying tolerances to TFM among fishes^3^. Detoxification of TFM is believed to occur through a combination of Phase I and Phase II biotransformation processes, which collectively act to enhance the water solubility of the compound to improve elimination^3^. It is generally believed that UDP-glucuronosyltransferase (Ugt) is the main enzyme responsible for TFM detoxification, via conjugation of TFM through the addition of a UDP-glucuronic acid^20–22^. The greater sensitivity of lampreys to TFM is believed to be related to lower Ugt activities compared to other fishes^20–, 22^, but the other cellular aspects of TFM detoxification are poorly characterized in fishes. Furthermore, interspecific variation in detoxification gene diversity is believed to be important in driving differences in toxicity tolerances among fishes^11^, which could suggest that available *ugt* genes impart differential TFM tolerances between sea lamprey and non-target fishes. Indeed, compared to zebrafish (*Danio rerio*; N =40), sea lamprey have far fewer *ugt* genes (N= 2) in their genome^23^. A few comparative studies have been conducted looking at sea lamprey and rainbow trout (*Oncorhynchus mykiss*), however little attention has been paid to the physiological effects of TFM exposure in other fishes^16–19^. As TFM detoxification primarily occurs in the liver, and the gills are one of the primary uptake sites, TFM physiological studies often focus on these two tissue types in fishes^3^. However, a broader characterization of TFM toxicity, particularly the expression patterns of Phase I and Phase II enzyme genes, in fishes is necessary.

We used a comparative transcriptomics approach to evaluate the effects of TFM on sea lamprey and bluegill, species with the lowest and highest reported tolerances to TFM, respectively^19^. The purpose of this study was to: 1) characterize interspecific variation in detoxification gene families expressed in the transcriptome following TFM exposure, 2) assess the potential mechanisms of TFM detoxification common to both species, and 3) identify key taxon-specific differences in the physiological response to TFM between sea lamprey and a TFM-tolerant teleost fish. We predicted that bluegill would have a greater diversity of detoxification mechanisms alongside a large robust transcriptomic detoxification response that contributes to their greater tolerance to TFM relative to sea lamprey. We also predicted that the molecular signatures of TFM toxicity would become more evident with increasing exposure duration as detoxification systems become exhausted; effects that would be more pronounced in the sea lamprey. These results will help to identify the evolutionary and genetic underpinnings of pesticide tolerance, which may be used to inform improvements in invasive species control efforts across a diversity of contexts.

## Results

### Detoxification gene expression patterns

Sea lamprey and bluegill were exposed to TFM for 6, 12, and 24 h at their species-specific 24 h-LC10 (sea lamprey = 2.21 mg L^-1^, bluegill = 22.06 mg L^-1^) with gill and liver tissue being sampled at 6, 12, and 24 h of exposure^19^. The mRNA from the gills and liver were extracted and sequenced using a whole transcriptome approach for the identification of gene targets associated with TFM detoxification and gene ontology (GO) term enrichment. As variation in TFM tolerance is believed to stem from differences in biotransformation capacities in fishes^3, 21^, we filtered for Phase I-III biotransformation transcripts from each species’ transcriptome. Proteins listed as being involved with detoxification, xenobiotic removal, and/or organic compound breakdown were retained as these functions likely made them important in TFM detoxification. Briefly, Phase I biotransformation generally includes the hydrolysis of organic compounds while Phase II biotransformation involves conjugation of organics with both processes enhancing xenobiotic solubility for easier elimination. Phase III biotransformation typically involves facilitating the excretion of the xenobiotic^24^.

Several Phase I detoxification transcripts were affected by TFM treatment in both species. In sea lamprey, transcripts associated with Phase I detoxification processes were only differentially expressed in the gills and consisted mainly of cytochrome P450s (six unique *cyp*’s; Table 1). Notably, this included both *cyp1a1* (6 h) and *cyp1b1* (24 h) with a paraoxonase (*pon1*) being the only non-*cyp* transcript differentially expressed (Table 1). Most *cyp* transcripts were upregulated in response to TFM exposure at 6 or 24 h, but *pon1*, *cyp2j2*, and *cyp2d3* were downregulated at 24 h. Contrastingly, bluegill demonstrated a larger induction of Phase I detoxification transcripts (nine unique transcripts, 28 differentially expressed transcripts) in both tissues across all exposure durations (Table 1). In bluegill gill, *cyp1a1* and *cyp1b1* were upregulated across all timepoints (Table 1). In the bluegill liver, several *cyp* transcripts responded positively to TFM treatment across exposure durations (e.g., *cyp1a1*, *cyp1b1*, *cyp2c31*), but three transcripts (*cyp2b2*, *cyp2k1*, *cyp2k6*) were downregulated at 24 h (Table 1).

**Table 1:**
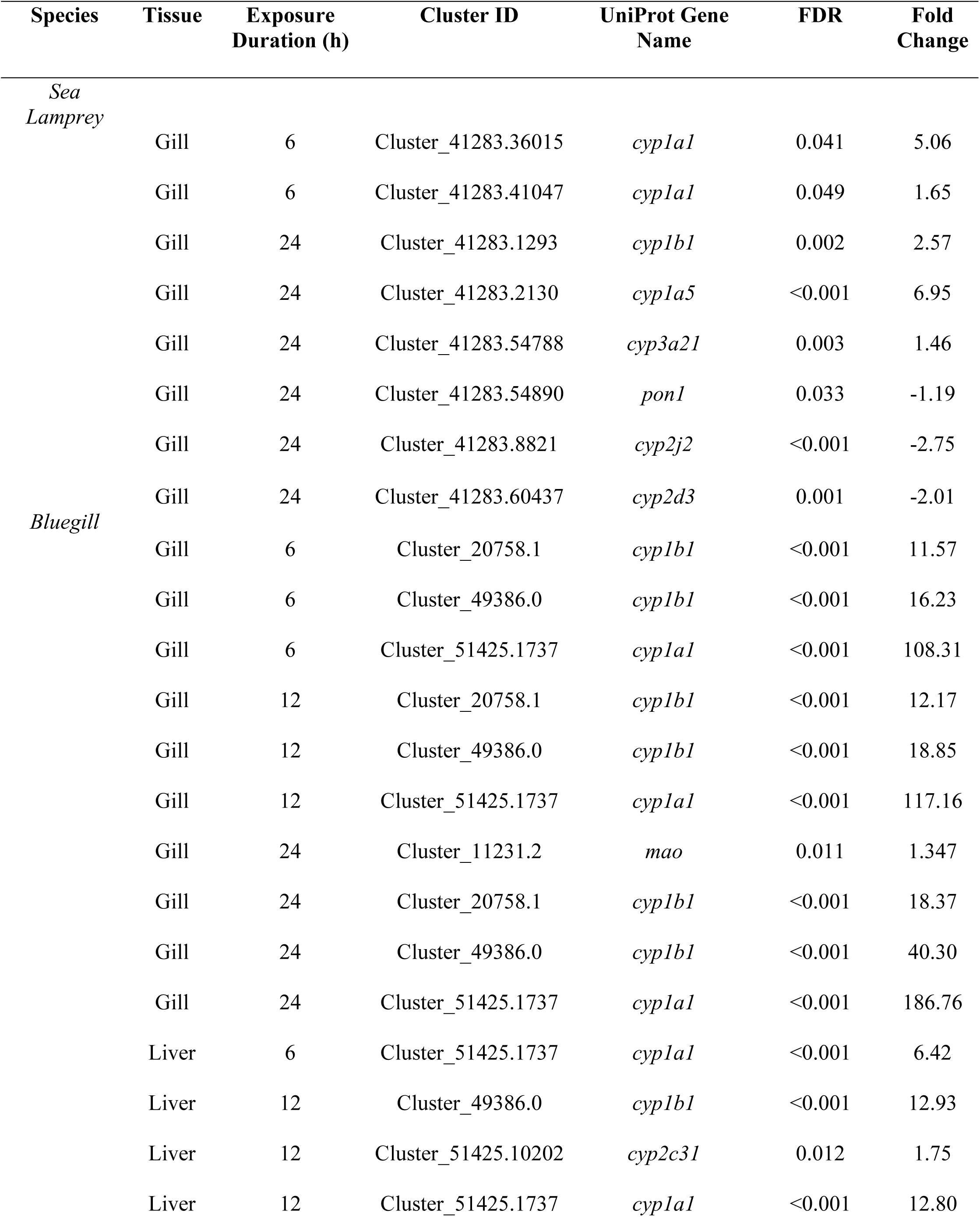
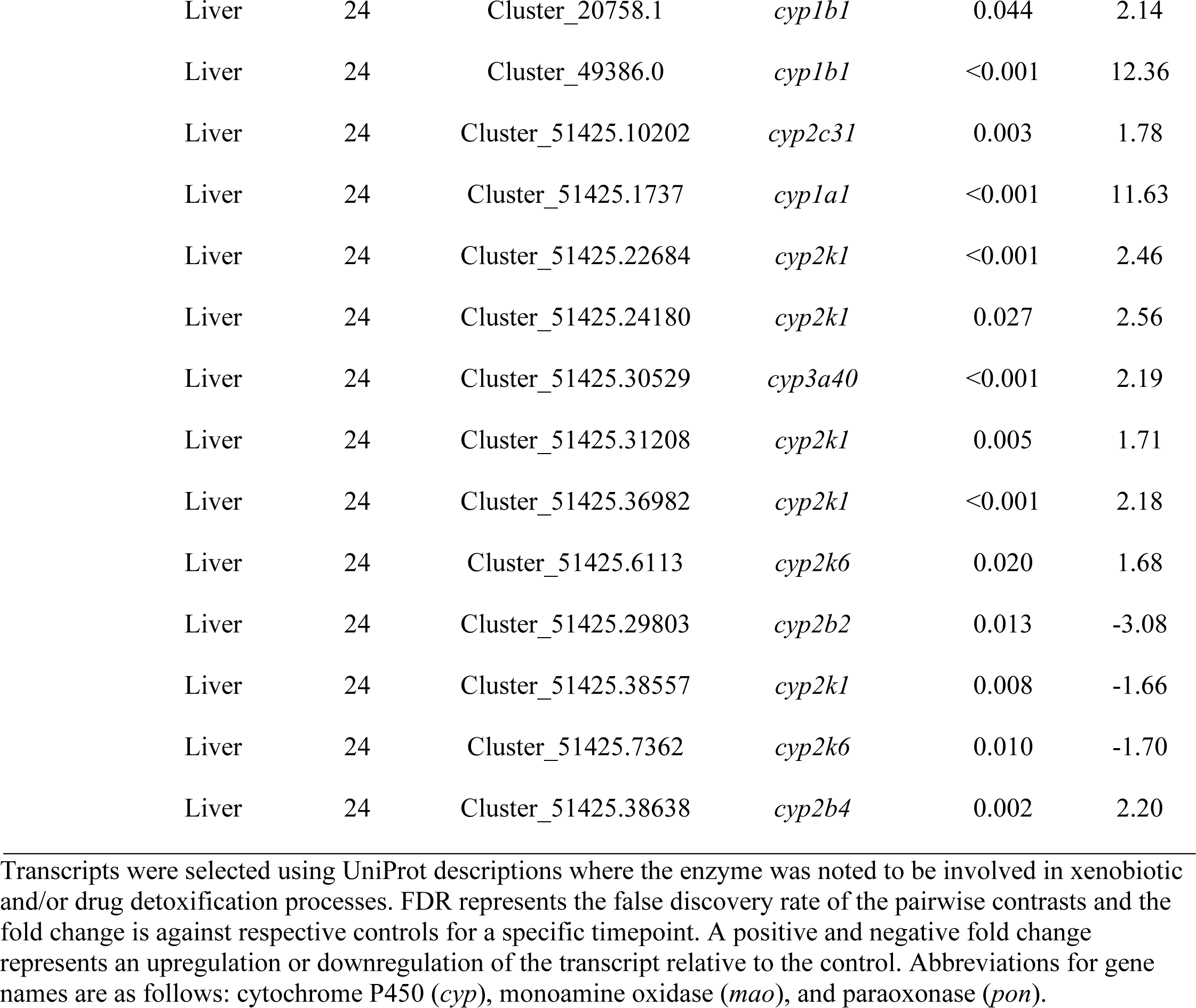
Differentially expressed transcripts associated with phase I detoxification for sea lamprey larvae (*Petromyzon marinus*) and bluegill (*Lepomis macrochirus*) following 6, 12, and 24 h TFM (3-trifluoromethyl-4’-nitrophenol) exposure.

| Species | Tissue | Exposure Duration (h) | Cluster ID | UniProt Gene Name | FDR | Fold Change |
| --- | --- | --- | --- | --- | --- | --- |
| *Sea Lamprey* | Gill | 6 | Cluster_41283.36015 | *cyp1a1* | 0.041 | 5.06 |
|  | Gill | 6 | Cluster_41283.41047 | *cyp1a1* | 0.049 | 1.65 |
|  | Gill | 24 | Cluster_41283.1293 | *cyp1b1* | 0.002 | 2.57 |
|  | Gill | 24 | Cluster_41283.2130 | *cyp1a5* | <0.001 | 6.95 |
|  | Gill | 24 | Cluster_41283.54788 | *cyp3a21* | 0.003 | 1.46 |
|  | Gill | 24 | Cluster_41283.54890 | *pon1* | 0.033 | -1.19 |
|  | Gill | 24 | Cluster_41283.8821 | *cyp2j2* | <0.001 | -2.75 |
|  | Gill | 24 | Cluster_41283.60437 | *cyp2d3* | 0.001 | -2.01 |
| *Bluegill* | Gill | 6 | Cluster_20758.1 | *cyp1b1* | <0.001 | 11.57 |
|  | Gill | 6 | Cluster_49386.0 | *cyp1b1* | <0.001 | 16.23 |
|  | Gill | 6 | Cluster_51425.1737 | *cyp1a1* | <0.001 | 108.31 |
|  | Gill | 12 | Cluster_20758.1 | *cyp1b1* | <0.001 | 12.17 |
|  | Gill | 12 | Cluster_49386.0 | *cyp1b1* | <0.001 | 18.85 |
|  | Gill | 12 | Cluster_51425.1737 | *cyp1a1* | <0.001 | 117.16 |
|  | Gill | 24 | Cluster_11231.2 | *mao* | 0.011 | 1.347 |
|  | Gill | 24 | Cluster_20758.1 | *cyp1b1* | <0.001 | 18.37 |
|  | Gill | 24 | Cluster_49386.0 | *cyp1b1* | <0.001 | 40.30 |
|  | Gill | 24 | Cluster_51425.1737 | *cyp1a1* | <0.001 | 186.76 |
|  | Liver | 6 | Cluster_51425.1737 | *cyp1a1* | <0.001 | 6.42 |
|  | Liver | 12 | Cluster_49386.0 | *cyp1b1* | <0.001 | 12.93 |
|  | Liver | 12 | Cluster_51425.10202 | *cyp2c31* | 0.012 | 1.75 |
|  | Liver | 12 | Cluster_51425.1737 | *cyp1a1* | <0.001 | 12.80 |

|  |  |  |  |  |  |
| --- | --- | --- | --- | --- | --- |
| Liver | 24 | Cluster_20758.1 | *cyp1b1* | 0.044 | 2.14 |
| Liver | 24 | Cluster_49386.0 | *cyp1b1* | <0.001 | 12.36 |
| Liver | 24 | Cluster_51425.10202 | *cyp2c31* | 0.003 | 1.78 |
| Liver | 24 | Cluster_51425.1737 | *cyp1a1* | <0.001 | 11.63 |
| Liver | 24 | Cluster_51425.22684 | *cyp2k1* | <0.001 | 2.46 |
| Liver | 24 | Cluster_51425.24180 | *cyp2k1* | 0.027 | 2.56 |
| Liver | 24 | Cluster_51425.30529 | *cyp3a40* | <0.001 | 2.19 |
| Liver | 24 | Cluster_51425.31208 | *cyp2k1* | 0.005 | 1.71 |
| Liver | 24 | Cluster_51425.36982 | *cyp2k1* | <0.001 | 2.18 |
| Liver | 24 | Cluster_51425.6113 | *cyp2k6* | 0.020 | 1.68 |
| Liver | 24 | Cluster_51425.29803 | *cyp2b2* | 0.013 | -3.08 |
| Liver | 24 | Cluster_51425.38557 | *cyp2k1* | 0.008 | -1.66 |
| Liver | 24 | Cluster_51425.7362 | *cyp2k6* | 0.010 | -1.70 |
| Liver | 24 | Cluster_51425.38638 | *cyp2b4* | 0.002 | 2.20 |

Transcripts were selected using UniProt descriptions where the enzyme was noted to be involved in xenobiotic and/or drug detoxification processes. FDR represents the false discovery rate of the pairwise contrasts and the fold change is against respective controls for a specific timepoint. A positive and negative fold change represents an upregulation or downregulation of the transcript relative to the control. Abbreviations for gene names are as follows: cytochrome P450 (*cyp*), monoamine oxidase (*mao*), and paraoxonase (*pon*).

In the transcriptomes of both species, we identified several *ugt*’s likely involved in the detoxification of TFM (Table 2). In sea lamprey, six clusters from the superTranscriptome mapped to *ugt*’s. In contrast, the bluegill transcriptome had 17 clusters which mapped to *ugt*’s (Table 2). There were also noticeable differences between sea lamprey and bluegill in expression of the *ugt* transcripts in response to TFM exposure. Treatment with TFM did not result in any *ugt* being differentially expressed in sea lamprey. Conversely, in bluegill, *ugt3* was upregulated in the gill across all three exposure durations (Table 3) and in the liver at 12 and 24 h of exposure. Interestingly, *ugt3* was not present in the lamprey transcriptome (Table 2). Hepatic *ugt2b9* transcript levels were also elevated under TFM exposure at 12 and 24 h in bluegill liver, and TFM exposure led to several *ugt* transcripts (i.e., *ugt1a2*, *ugt1a5*, *ugt2a1*) being downregulated in the liver of bluegill at 12 and 24 h of TFM exposure (Table 3).

**Table 2:**
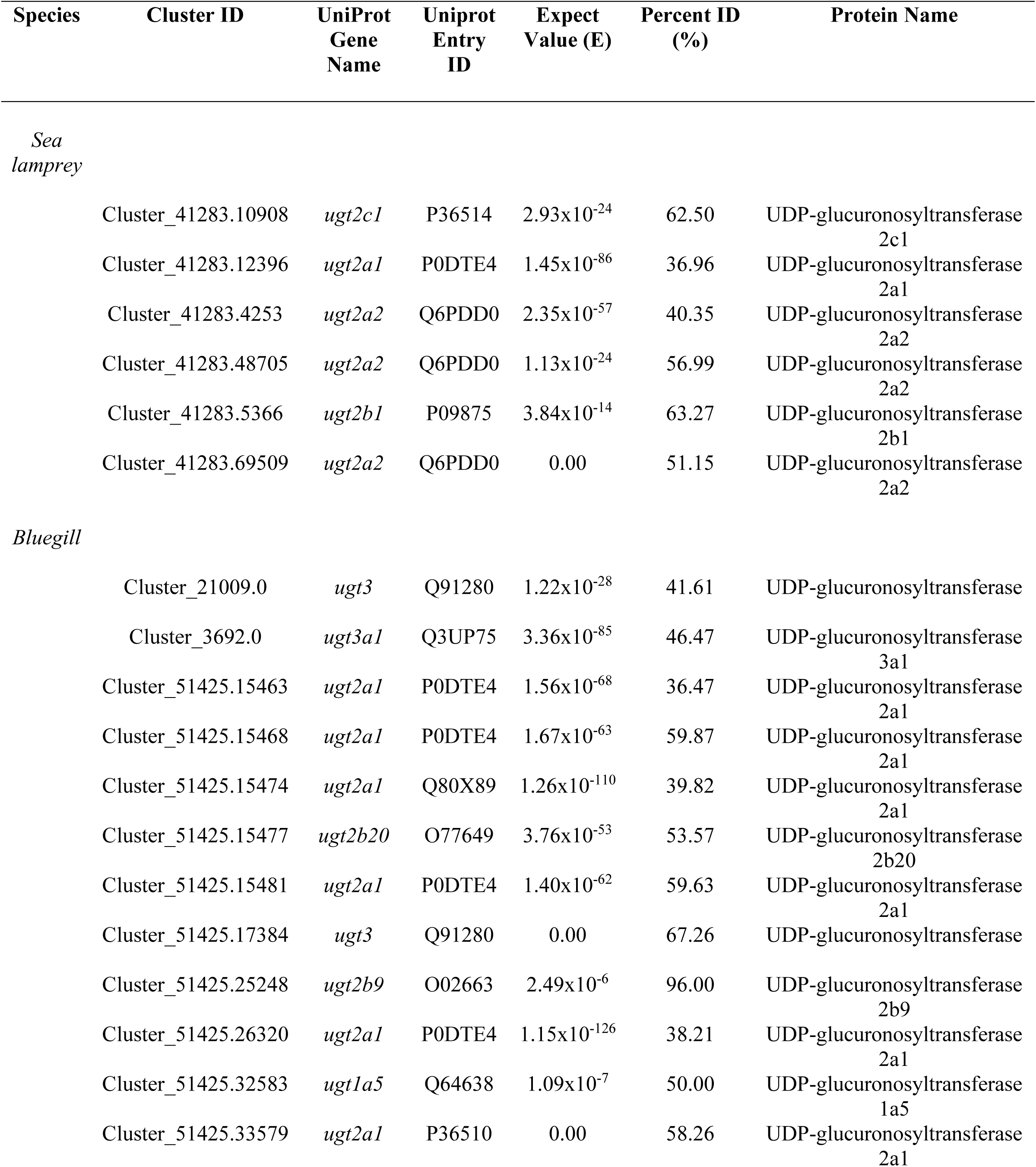
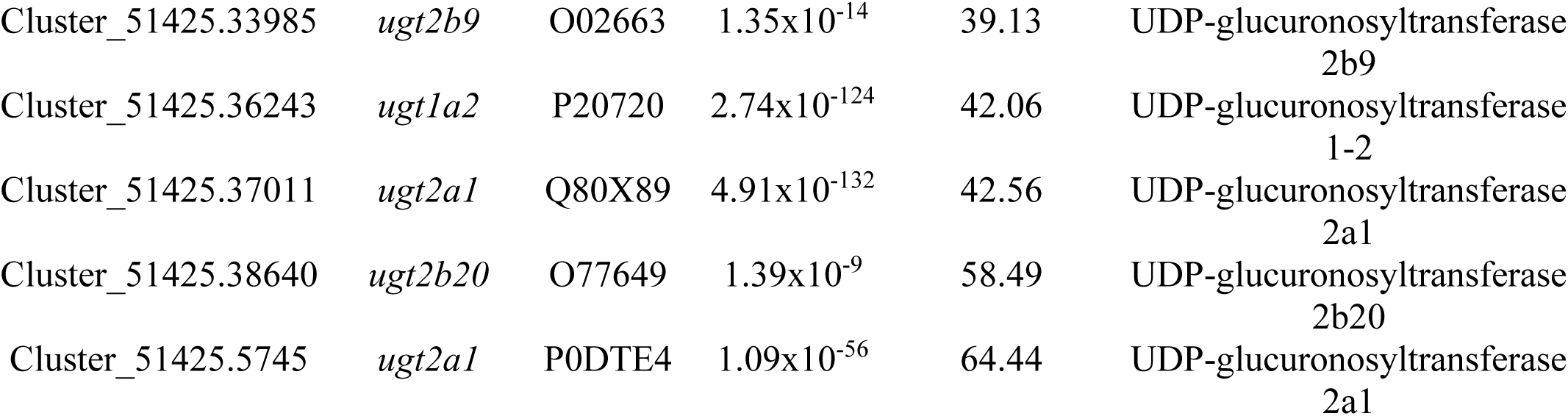
Identification of all UDP-glucuronosyltransferase (*ugt*) transcripts in the annotated transcriptomes of sea lamprey larvae (*Petromyzon marinus*) and bluegill (*Lepomis macrochirus*).

| Species | Cluster ID | UniProt Gene Name | Uniprot Entry ID | Expect Value (E) | Percent ID (%) | Protein Name |
| --- | --- | --- | --- | --- | --- | --- |
| *Sea lamprey* |  |  |  |  |  |  |
|  | Cluster_41283.10908 | *ugt2c1* | P36514 | 2.93x10<sup>-24</sup> | 62.50 | UDP-glucuronosyltransferase 2c1 |
|  | Cluster_41283.12396 | *ugt2a1* | P0DTE4 | 1.45x10<sup>-86</sup> | 36.96 | UDP-glucuronosyltransferase 2a1 |
|  | Cluster_41283.4253 | *ugt2a2* | Q6PDD0 | 2.35x10<sup>-57</sup> | 40.35 | UDP-glucuronosyltransferase 2a2 |
|  | Cluster_41283.48705 | *ugt2a2* | Q6PDD0 | 1.13x10<sup>-24</sup> | 56.99 | UDP-glucuronosyltransferase 2a2 |
|  | Cluster_41283.5366 | *ugt2b1* | P09875 | 3.84x10<sup>-14</sup> | 63.27 | UDP-glucuronosyltransferase 2b1 |
|  | Cluster_41283.69509 | *ugt2a2* | Q6PDD0 | 0.00 | 51.15 | UDP-glucuronosyltransferase 2a2 |
| *Bluegill* |  |  |  |  |  |  |
|  | Cluster_21009.0 | *ugt3* | Q91280 | 1.22x10<sup>-28</sup> | 41.61 | UDP-glucuronosyltransferase |
|  | Cluster_3692.0 | *ugt3a1* | Q3UP75 | 3.36x10<sup>-85</sup> | 46.47 | UDP-glucuronosyltransferase 3a1 |
|  | Cluster_51425.15463 | *ugt2a1* | P0DTE4 | 1.56x10<sup>-68</sup> | 36.47 | UDP-glucuronosyltransferase 2a1 |
|  | Cluster_51425.15468 | *ugt2a1* | P0DTE4 | 1.67x10<sup>-63</sup> | 59.87 | UDP-glucuronosyltransferase 2a1 |
|  | Cluster_51425.15474 | *ugt2a1* | Q80X89 | 1.26x10<sup>-110</sup> | 39.82 | UDP-glucuronosyltransferase 2a1 |
|  | Cluster_51425.15477 | *ugt2b20* | O77649 | 3.76x10<sup>-53</sup> | 53.57 | UDP-glucuronosyltransferase 2b20 |
|  | Cluster_51425.15481 | *ugt2a1* | P0DTE4 | 1.40x10<sup>-62</sup> | 59.63 | UDP-glucuronosyltransferase 2a1 |
|  | Cluster_51425.17384 | *ugt3* | Q91280 | 0.00 | 67.26 | UDP-glucuronosyltransferase |
|  | Cluster_51425.25248 | *ugt2b9* | O02663 | 2.49x10<sup>-6</sup> | 96.00 | UDP-glucuronosyltransferase 2b9 |
|  | Cluster_51425.26320 | *ugt2a1* | P0DTE4 | 1.15x10<sup>-126</sup> | 38.21 | UDP-glucuronosyltransferase 2a1 |
|  | Cluster_51425.32583 | *ugt1a5* | Q64638 | 1.09x10<sup>-7</sup> | 50.00 | UDP-glucuronosyltransferase 1a5 |
|  | Cluster_51425.33579 | *ugt2a1* | P36510 | 0.00 | 58.26 | UDP-glucuronosyltransferase 2a1 |

|  |  |  |  |  |  |
| --- | --- | --- | --- | --- | --- |
| Cluster_51425.33985 | *ugt2b9* | O02663 | $1.35 \times 10^{-14}$ | 39.13 | UDP-glucuronosyltransferase<br>2b9 |
| Cluster_51425.36243 | *ugt1a2* | P20720 | $2.74 \times 10^{-124}$ | 42.06 | UDP-glucuronosyltransferase<br>1-2 |
| Cluster_51425.37011 | *ugt2a1* | Q80X89 | $4.91 \times 10^{-132}$ | 42.56 | UDP-glucuronosyltransferase<br>2a1 |
| Cluster_51425.38640 | *ugt2b20* | O77649 | $1.39 \times 10^{-9}$ | 58.49 | UDP-glucuronosyltransferase<br>2b20 |
| Cluster_51425.5745 | *ugt2a1* | P0DTE4 | $1.09 \times 10^{-56}$ | 64.44 | UDP-glucuronosyltransferase<br>2a1 |

**Table 3:**
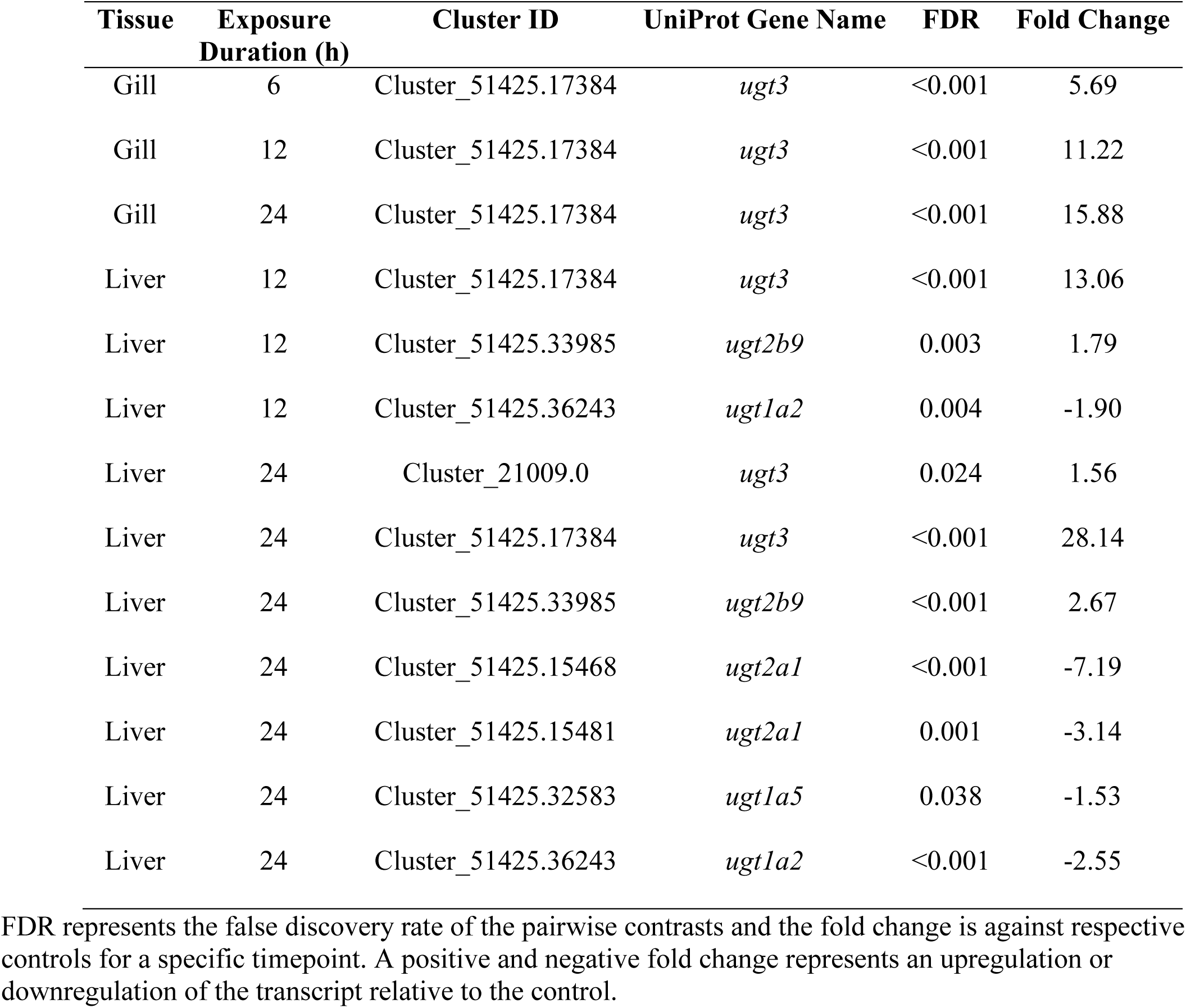
Summary of the differentially expressed UDP-glucuronosyltransferase (UGT) transcripts for bluegill (*Lepomis macrochirus*) following 6, 12, and 24 h TFM (3-trifluoromethyl-4’-nitrophenol) exposure.

| Tissue | Exposure Duration (h) | Cluster ID | UniProt Gene Name | FDR | Fold Change |
| --- | --- | --- | --- | --- | --- |
| Gill | 6 | Cluster_51425.17384 | *ugt3* | <0.001 | 5.69 |
| Gill | 12 | Cluster_51425.17384 | *ugt3* | <0.001 | 11.22 |
| Gill | 24 | Cluster_51425.17384 | *ugt3* | <0.001 | 15.88 |
| Liver | 12 | Cluster_51425.17384 | *ugt3* | <0.001 | 13.06 |
| Liver | 12 | Cluster_51425.33985 | *ugt2b9* | 0.003 | 1.79 |
| Liver | 12 | Cluster_51425.36243 | *ugt1a2* | 0.004 | -1.90 |
| Liver | 24 | Cluster_21009.0 | *ugt3* | 0.024 | 1.56 |
| Liver | 24 | Cluster_51425.17384 | *ugt3* | <0.001 | 28.14 |
| Liver | 24 | Cluster_51425.33985 | *ugt2b9* | <0.001 | 2.67 |
| Liver | 24 | Cluster_51425.15468 | *ugt2a1* | <0.001 | -7.19 |
| Liver | 24 | Cluster_51425.15481 | *ugt2a1* | 0.001 | -3.14 |
| Liver | 24 | Cluster_51425.32583 | *ugt1a5* | 0.038 | -1.53 |
| Liver | 24 | Cluster_51425.36243 | *ugt1a2* | <0.001 | -2.55 |

FDR represents the false discovery rate of the pairwise contrasts and the fold change is against respective controls for a specific timepoint. A positive and negative fold change represents an upregulation or downregulation of the transcript relative to the control.

There was also a noticeable difference between bluegill and sea lamprey in the expression patterns for other transcripts involved in Phase II detoxification (Table 4). In sea lamprey, there were only three differentially expressed transcripts for Phase II genes, including glutathione S-transferase (*gstt3*), sulfotransferase (*sult6b1*), and nicotinamide N-methyltransferase (*nnmt*), in both liver and gill, all at 24 h of exposure (Table 4). In bluegill, several sulfotransferases and a single N-acetyltransferase (*nat8*; Table 4) transcripts were upregulated in the gill. In the bluegill liver, Phase II detoxification transcripts were only differentially regulated at 24 h of TFM exposure with N-acetyltransferases (*nat2* and *nat8*) being the only upregulated transcripts and two glutathione S-transferases (*gstt3* and *gstt1*), a thiopurine S-methyltransferase (*tpmt*), and a sulfotransferase (*sult1c1*) being downregulated (Table 4).

**Table 4:**
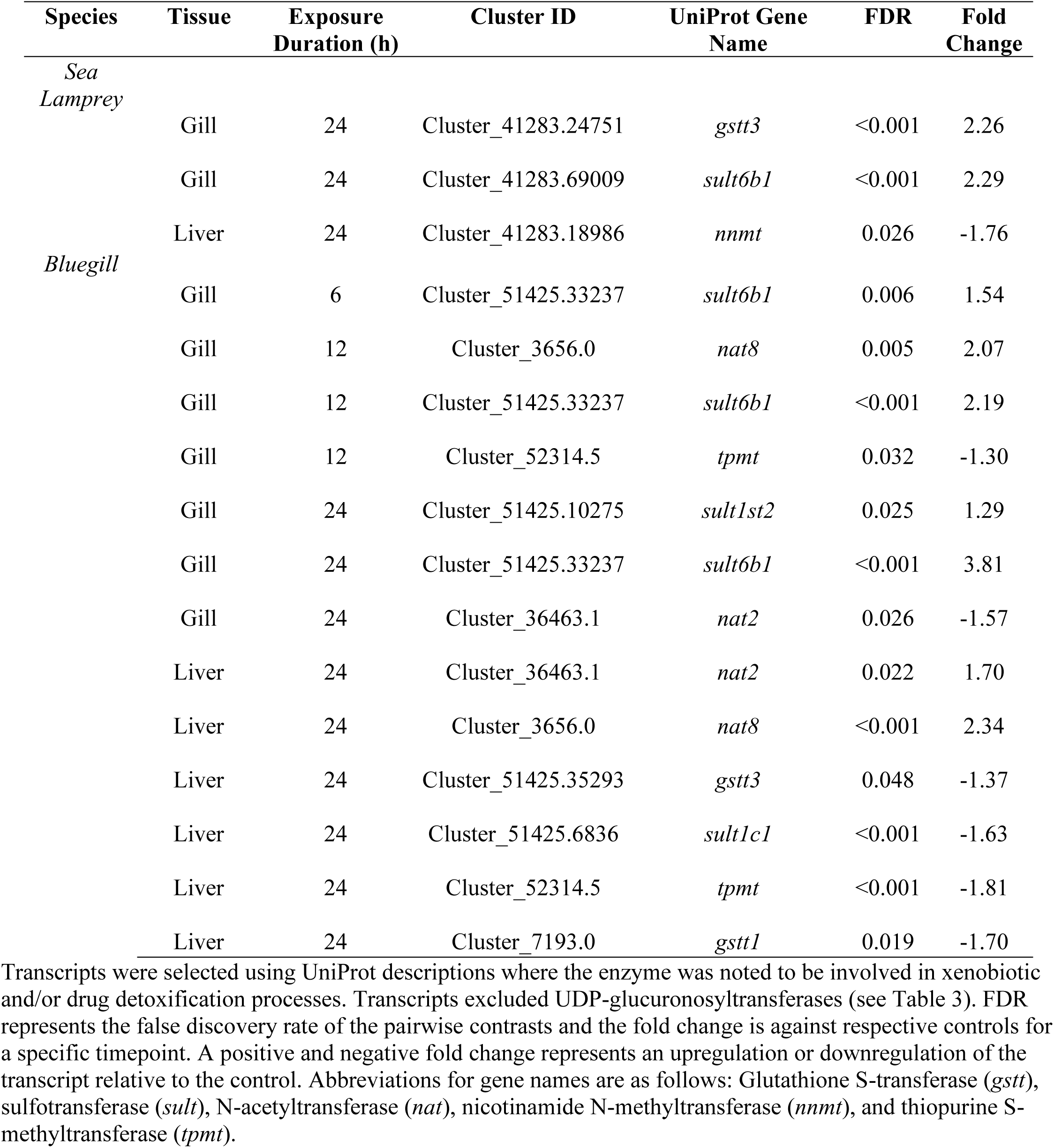
Differentially expressed transcripts associated with phase II detoxification for sea lamprey larvae (*Petromyzon marinus*) and bluegill (*Lepomis macrochirus*) following 6, 12, and 24 h TFM (3-trifluoromethyl-4’-nitrophenol) exposure.

| Species | Tissue | Exposure<br>Duration (h) | Cluster ID | UniProt Gene<br>Name | FDR | Fold<br>Change |
| --- | --- | --- | --- | --- | --- | --- |
| *Sea*<br>*Lamprey* | Gill | 24 | Cluster_41283.24751 | *gstt3* | <0.001 | 2.26 |
|  | Gill | 24 | Cluster_41283.69009 | *sult6b1* | <0.001 | 2.29 |
|  | Liver | 24 | Cluster_41283.18986 | *nnmt* | 0.026 | -1.76 |
| *Bluegill* | Gill | 6 | Cluster_51425.33237 | *sult6b1* | 0.006 | 1.54 |
|  | Gill | 12 | Cluster_3656.0 | *nat8* | 0.005 | 2.07 |
|  | Gill | 12 | Cluster_51425.33237 | *sult6b1* | <0.001 | 2.19 |
|  | Gill | 12 | Cluster_52314.5 | *tpmt* | 0.032 | -1.30 |
|  | Gill | 24 | Cluster_51425.10275 | *sult1st2* | 0.025 | 1.29 |
|  | Gill | 24 | Cluster_51425.33237 | *sult6b1* | <0.001 | 3.81 |
|  | Gill | 24 | Cluster_36463.1 | *nat2* | 0.026 | -1.57 |
|  | Liver | 24 | Cluster_36463.1 | *nat2* | 0.022 | 1.70 |
|  | Liver | 24 | Cluster_3656.0 | *nat8* | <0.001 | 2.34 |
|  | Liver | 24 | Cluster_51425.35293 | *gstt3* | 0.048 | -1.37 |
|  | Liver | 24 | Cluster_51425.6836 | *sult1c1* | <0.001 | -1.63 |
|  | Liver | 24 | Cluster_52314.5 | *tpmt* | <0.001 | -1.81 |
|  | Liver | 24 | Cluster_7193.0 | *gstt1* | 0.019 | -1.70 |

Differential expression of Phase III detoxification transcripts was limited to 24 h of exposure in sea lamprey. In the gills, five ATP-binding cassettes (*abc*; *abcc3*, *abccc5*, *abcb1*, *abcb11*, and *abcg2*) were upregulated in response to TFM, whereas the two solute carrier family transcripts (*slc*; *slc35d2* and *slc22a15b*) were downregulated (Table 5). In the lamprey liver, a single *abc* and *slc* were the only transcripts differentially expressed in response to TFM (Table 5). In contrast, the gills of bluegill exhibited differential expression of Phase III transcripts across all three timepoints consisting of four unique *abc*’s (*abca3*, *abcb1*, *abcb6*, and *abcc4*) and nine unique *slc*’s (Table 5). In most instances, these branchial *abc*’s and *slc*’s were upregulated in bluegill. In the bluegill liver, a single *abc* (*abcb6*) was found to be upregulated at 12 and 24 h of TFM exposure (Table 5). Aside from *slc47a1*, all of the five-remaining hepatic *slc*’s were found be upregulated under TFM treatment and with differential expression occurring at 12 and 24 h of exposure (Table 5).

**Table 5:**
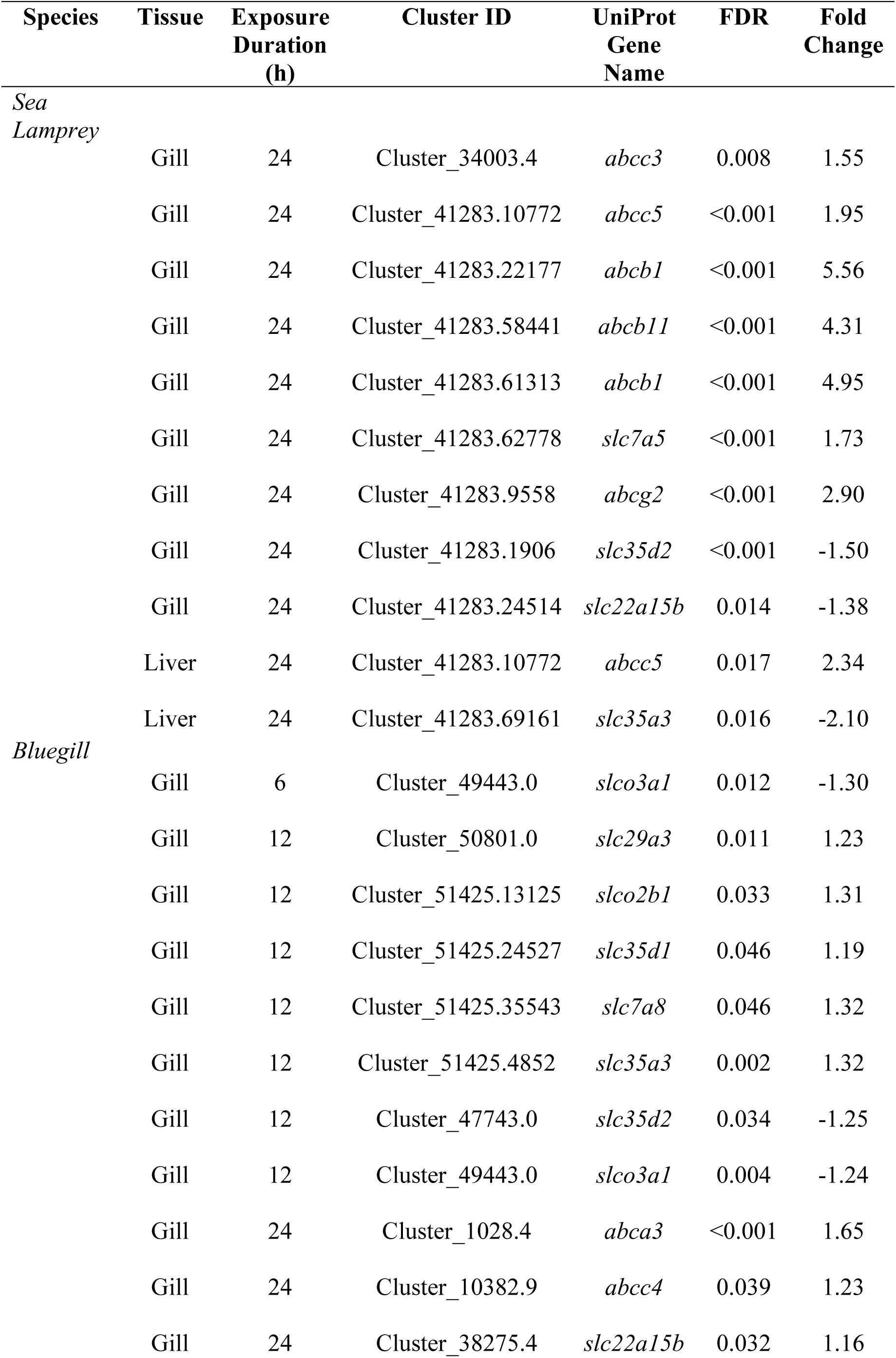
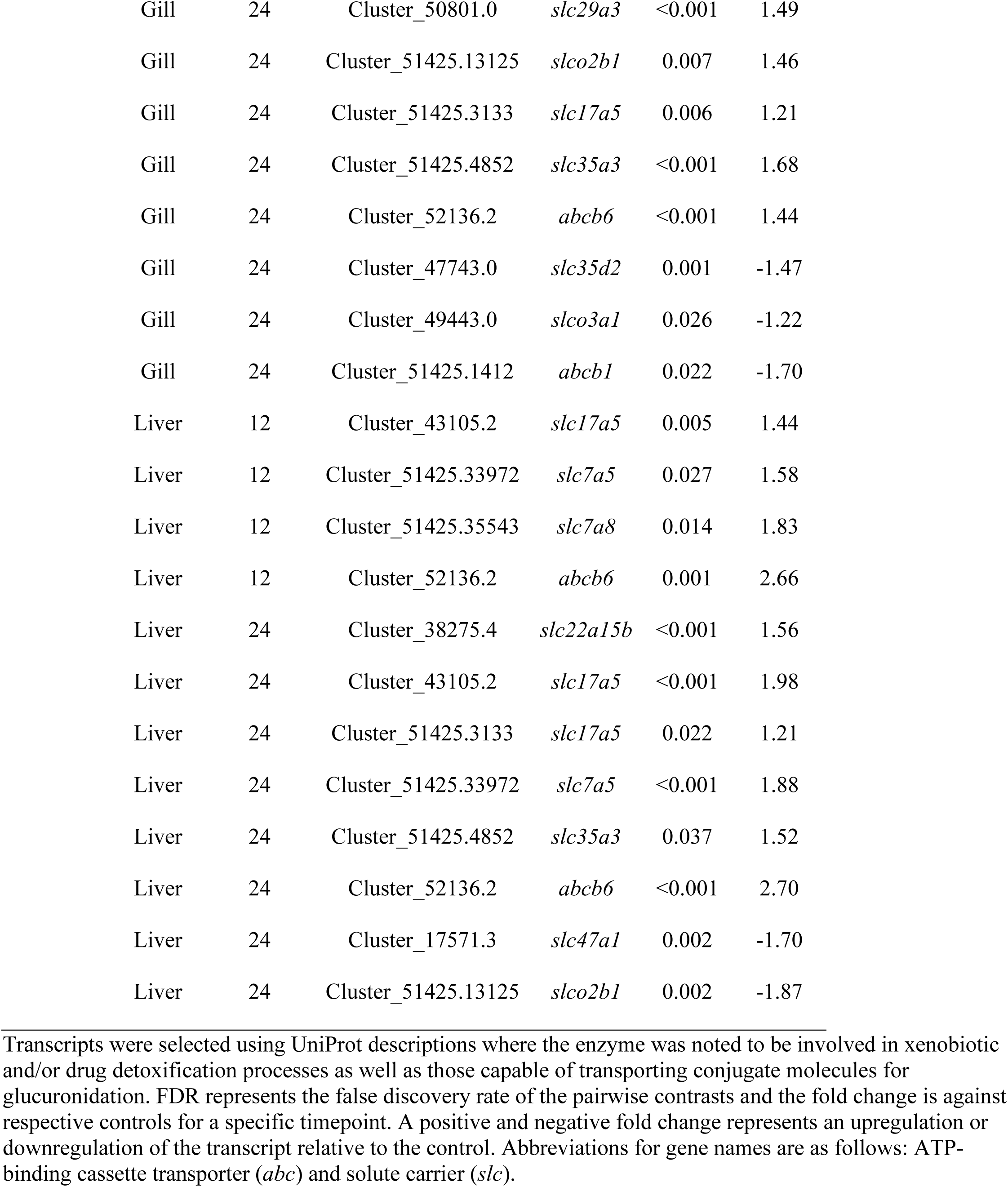
Differentially expressed transcripts associated with phase III detoxification for sea lamprey larvae (*Petromyzon marinus*) and bluegill (*Lepomis macrochirus*) following 6, 12, and 24 h TFM (3-trifluoromethyl-4’-nitrophenol) exposure.

| Species | Tissue | Exposure Duration (h) | Cluster ID | UniProt Gene Name | FDR | Fold Change |
| --- | --- | --- | --- | --- | --- | --- |
| *Sea Lamprey* | Gill | 24 | Cluster_34003.4 | *abcc3* | 0.008 | 1.55 |
|  | Gill | 24 | Cluster_41283.10772 | *abcc5* | <0.001 | 1.95 |
|  | Gill | 24 | Cluster_41283.22177 | *abcb1* | <0.001 | 5.56 |
|  | Gill | 24 | Cluster_41283.58441 | *abcb11* | <0.001 | 4.31 |
|  | Gill | 24 | Cluster_41283.61313 | *abcb1* | <0.001 | 4.95 |
|  | Gill | 24 | Cluster_41283.62778 | *slc7a5* | <0.001 | 1.73 |
|  | Gill | 24 | Cluster_41283.9558 | *abcg2* | <0.001 | 2.90 |
|  | Gill | 24 | Cluster_41283.1906 | *slc35d2* | <0.001 | -1.50 |
|  | Gill | 24 | Cluster_41283.24514 | *slc22a15b* | 0.014 | -1.38 |
|  | Liver | 24 | Cluster_41283.10772 | *abcc5* | 0.017 | 2.34 |
|  | Liver | 24 | Cluster_41283.69161 | *slc35a3* | 0.016 | -2.10 |
| *Bluegill* | Gill | 6 | Cluster_49443.0 | *slco3a1* | 0.012 | -1.30 |
|  | Gill | 12 | Cluster_50801.0 | *slc29a3* | 0.011 | 1.23 |
|  | Gill | 12 | Cluster_51425.13125 | *slco2b1* | 0.033 | 1.31 |
|  | Gill | 12 | Cluster_51425.24527 | *slc35d1* | 0.046 | 1.19 |
|  | Gill | 12 | Cluster_51425.35543 | *slc7a8* | 0.046 | 1.32 |
|  | Gill | 12 | Cluster_51425.4852 | *slc35a3* | 0.002 | 1.32 |
|  | Gill | 12 | Cluster_47743.0 | *slc35d2* | 0.034 | -1.25 |
|  | Gill | 12 | Cluster_49443.0 | *slco3a1* | 0.004 | -1.24 |
|  | Gill | 24 | Cluster_1028.4 | *abca3* | <0.001 | 1.65 |
|  | Gill | 24 | Cluster_10382.9 | *abcc4* | 0.039 | 1.23 |
|  | Gill | 24 | Cluster_38275.4 | *slc22a15b* | 0.032 | 1.16 |

|  |  |  |  |  |  |
| --- | --- | --- | --- | --- | --- |
| Gill | 24 | Cluster_50801.0 | *slc29a3* | <0.001 | 1.49 |
| Gill | 24 | Cluster_51425.13125 | *slco2b1* | 0.007 | 1.46 |
| Gill | 24 | Cluster_51425.3133 | *slc17a5* | 0.006 | 1.21 |
| Gill | 24 | Cluster_51425.4852 | *slc35a3* | <0.001 | 1.68 |
| Gill | 24 | Cluster_52136.2 | *abcb6* | <0.001 | 1.44 |
| Gill | 24 | Cluster_47743.0 | *slc35d2* | 0.001 | -1.47 |
| Gill | 24 | Cluster_49443.0 | *slco3a1* | 0.026 | -1.22 |
| Gill | 24 | Cluster_51425.1412 | *abcb1* | 0.022 | -1.70 |
| Liver | 12 | Cluster_43105.2 | *slc17a5* | 0.005 | 1.44 |
| Liver | 12 | Cluster_51425.33972 | *slc7a5* | 0.027 | 1.58 |
| Liver | 12 | Cluster_51425.35543 | *slc7a8* | 0.014 | 1.83 |
| Liver | 12 | Cluster_52136.2 | *abcb6* | 0.001 | 2.66 |
| Liver | 24 | Cluster_38275.4 | *slc22a15b* | <0.001 | 1.56 |
| Liver | 24 | Cluster_43105.2 | *slc17a5* | <0.001 | 1.98 |
| Liver | 24 | Cluster_51425.3133 | *slc17a5* | 0.022 | 1.21 |
| Liver | 24 | Cluster_51425.33972 | *slc7a5* | <0.001 | 1.88 |
| Liver | 24 | Cluster_51425.4852 | *slc35a3* | 0.037 | 1.52 |
| Liver | 24 | Cluster_52136.2 | *abcb6* | <0.001 | 2.70 |
| Liver | 24 | Cluster_17571.3 | *slc47a1* | 0.002 | -1.70 |
| Liver | 24 | Cluster_51425.13125 | *slco2b1* | 0.002 | -1.87 |

Transcripts were selected using UniProt descriptions where the enzyme was noted to be involved in xenobiotic and/or drug detoxification processes as well as those capable of transporting conjugate molecules for glucuronidation. FDR represents the false discovery rate of the pairwise contrasts and the fold change is against respective controls for a specific timepoint. A positive and negative fold change represents an upregulation or downregulation of the transcript relative to the control. Abbreviations for gene names are as follows: ATP-binding cassette transporter (*abc*) and solute carrier (*slc*).

### Overall transcriptome patterns

The overall number of differentially expressed transcripts and their timing were different in the two species. In the sea lamprey, there was a delayed transcriptional response in the gills following TFM exposure (Fig. 1a). At 6 and 12 h, we observed only 139 and 9 differentially expressed superTranscriptome clusters, respectively (hereafter referred to as ‘clusters’). However, by 24 h, there were 7,055 differentially expressed clusters. In contrast, bluegill gill expression patterns were marked by a stepwise increase in the number of differentially expressed superTranscriptome clusters. A total of 595 and 2,662 differentially expressed clusters were already observed at 6 and 12 h of TFM exposure, respectively, although the total at 24 h (4,370) was less than the 24-h total in sea lamprey gills (Fig. 1b).

**Figure 1:**
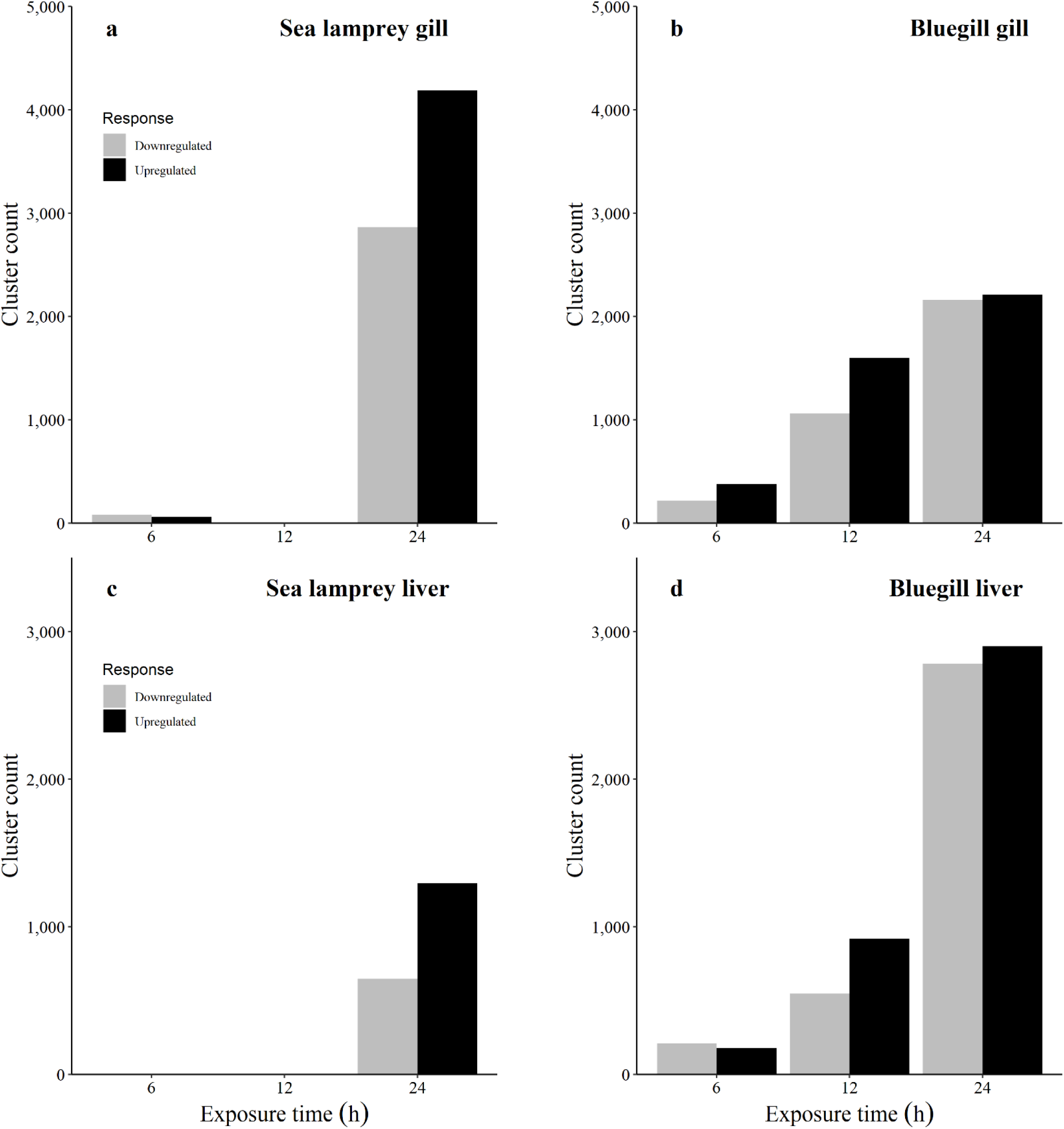
Total counts of differentially expressed superTranscriptome clusters in the gills (a,b) and liver (c,d) of sea lamprey larvae and bluegill following a 24 h TFM exposure. Grey bars represent differentially expressed clusters that were downregulated whereas black bars represent upregulated clusters. All clusters were deemed to be significant at a false discovery rate (FDR) of 29 α = 0.05.

The livers of sea lamprey showed a muted pattern of transcriptional activity under TFM exposure with no changes in transcription within the first 12 h of exposure, and only 1,940 differentially expressed clusters by 24 h (Fig. 1c). In bluegill, there were moderate levels of differential expression in the liver within the first 12 h of TFM exposure (388 and 1,464 at 6 and 12 h, respectively), and the number had increased to 5,685 differentially expressed clusters by 24 h of exposure (Fig. 1d).

### Lamprey GO term enrichment

In sea lamprey gills, TFM treatment resulted in an enrichment of GO terms associated with cellular growth and proliferation, immune function, and metabolism by 24 h of exposure. Specifically, in transcripts that were upregulated, GO terms associated with cellular proliferation and growth included regulation of cell proliferation, Ras protein signaling, regulation of I-κB kinase/signaling, and regulation of the cell cycle (Fig. 2a). GO terms related to immune function including viral life cycle, antigen processing, and negative regulation of type I interferon were also enriched for upregulated transcripts in sea lamprey (Fig. 2a). An upregulation in GO terms associated with energy metabolism, including “cellular response to insulin stimulus”, and “regulation of gluconeogenesis”, also featured prominently in the sea lamprey gill (Fig. 2a). Of the gluconeogenic transcripts that were enriched, we specifically observed an upregulation in mitochondrial pyruvate dehydrogenase kinase 2 (*pdk2*) and glycerol-3-phosphate phosphatase (*pgp*).

**Figure 2:**
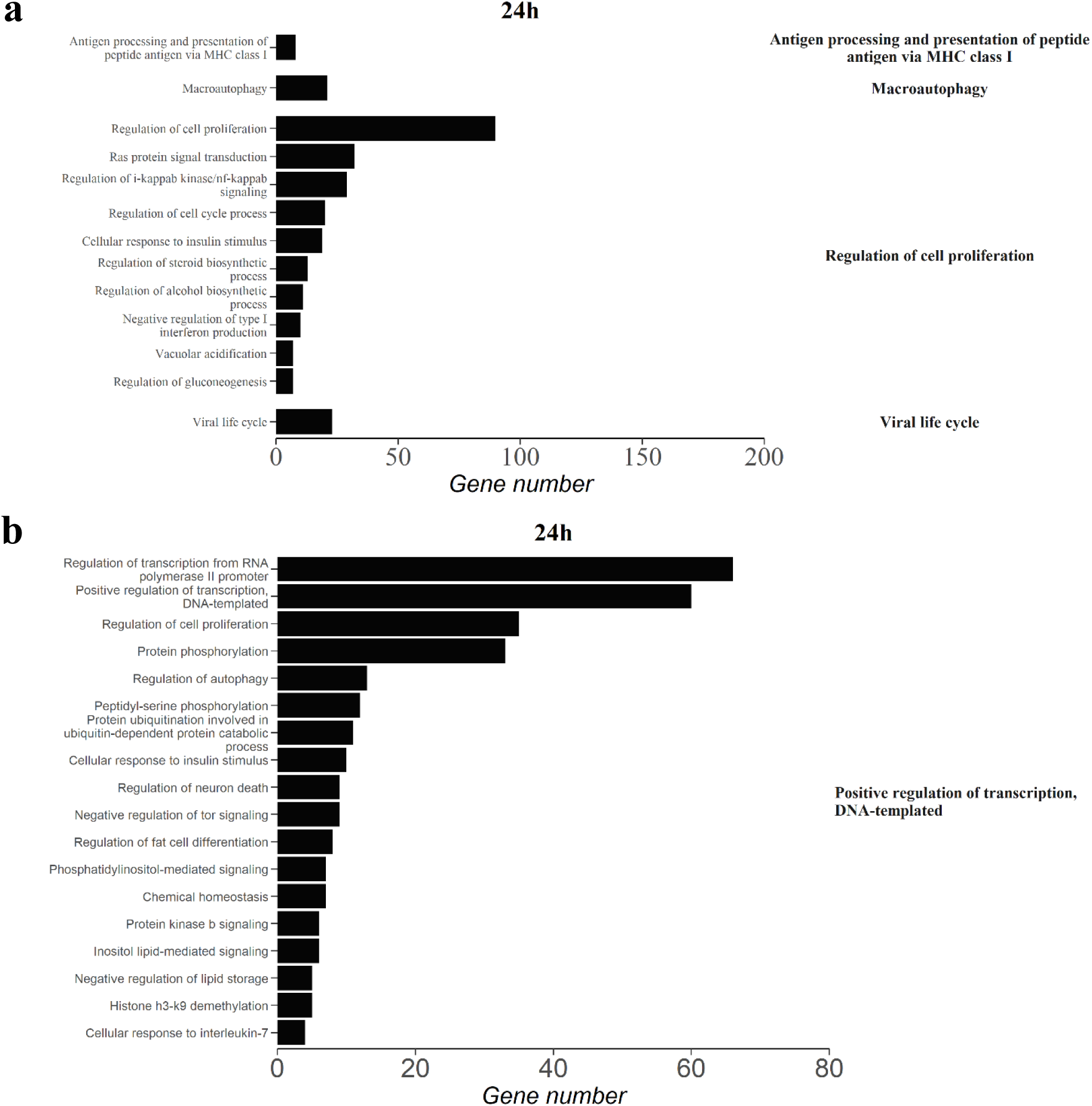
Summary of enriched gene ontology (GO) terms with transcripts that were upregulated following 24 h of TFM (3-trifluoromethyl-4’-nitrophenol; 2.21 mg L^-1^ nominal) exposure in the gills (**a**) and liver (**b**) of sea lamprey (*Petromyzon marinus*) larvae. Transcripts were considered differentially regulated at a false discovery rate < 0.05. Only GO terms from the functional analysis with an adjusted p < 0.05 and at least four transcripts were considered significantly enriched. REVIGO was used to summarize GO terms to reduce redundancy and group according to similarity (right labels).

In sea lamprey liver, there were no differentially expressed transcripts at 6 h and 12 h of TFM exposure (Fig. 1c). At 24 h, there was an enrichment of upregulated transcripts associated with the GO term “positive regulation of transcription” (Fig. 2b). Within this parent term, cell growth and survival GO terms were enriched, which included regulation of cell proliferation, fat cell differentiation, autophagy, and ubiquitination. Additionally, processes related to energy metabolism appeared to be upregulated as there was enrichment of transcripts associated with “cellular response to insulin stimulus” (e.g., *igf1r*, *insr*, *pck2*, *pdk2*) and “negative regulation of lipid storage” (e.g., *abca1*, *nfkbia*, *osbpl8*; Fig. 2b).

### Bluegill GO term enrichment

In bluegill gills, there was enrichment of upregulated transcripts associated with cellular growth, proliferation, and death. For example, at 6 h of TFM exposure, there was enrichment of apoptotic processes regulation (Fig. 3a). By 12 h, this included a “positive regulation of cell proliferation”, “transforming growth factor β (TGF-β) responsiveness”, “regulation of autophagy”, “regulation of apoptotic process”, and “ubiquitin dependent processes” (Fig. 3a). By 24 h, enrichment was largely restricted to cellular death including GO terms related to apoptosis and macroautophagy (Fig. 3a). Several transcripts associated with apoptosis, including *akt2*, several *bmp* and *map3k* transcripts, and *casp3*, were upregulated under TFM. For downregulated transcripts in the gill, there was also further enrichment of processes related to cellular growth and death (Fig. 3b). At 6 h, we observed a negative regulation of several growth-related processes including a “positive regulation of cell proliferation” and “Wnt signaling” (Fig. 3b).

**Figure 3:**
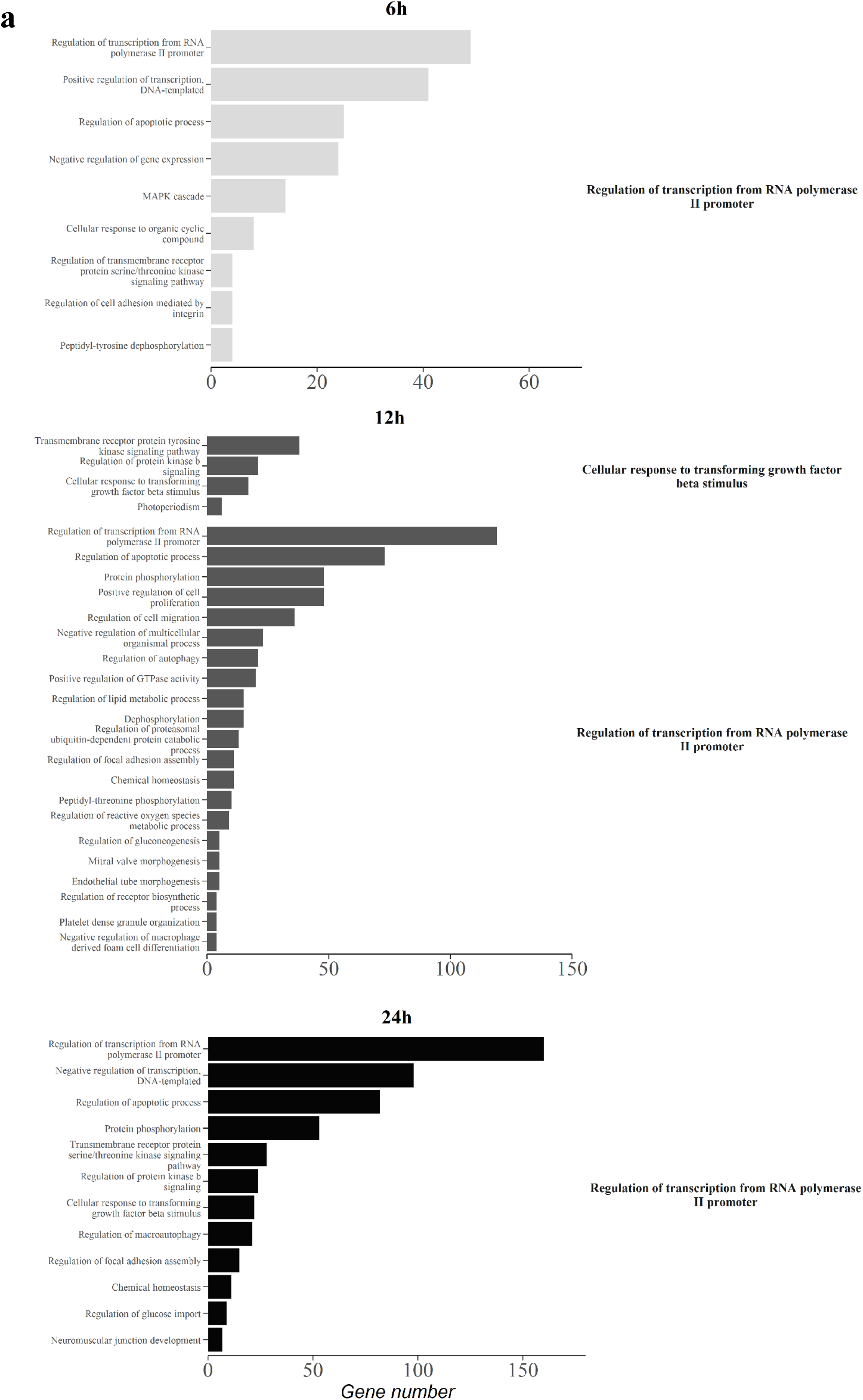
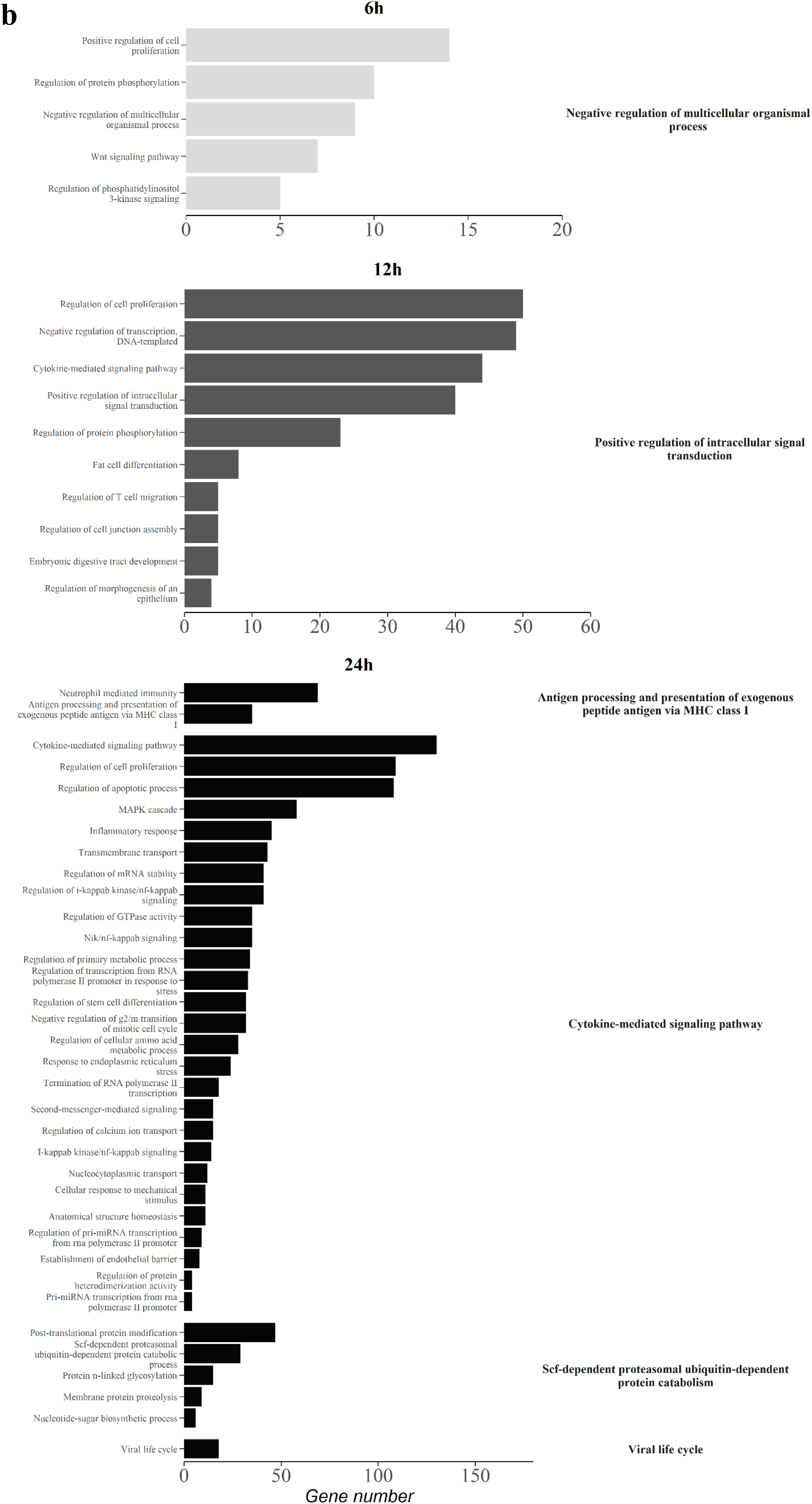
Summary of enriched gene ontology (GO) terms with transcripts that were upregulated (**a**) and downregulated (**b**) following 6 h (light grey), 12 h (dark grey), and 24 h (black) of TFM (3-trifluoromethyl-4’-nitrophenol; 22.06 mg L^-1^ nominal) exposure in the gills of bluegill (*Lepomis macrochirus*). Transcripts were considered differentially regulated at a false discovery rate < 0.05. Only GO terms from the functional analysis with an adjusted p < 0.05 and at least four transcripts were considered significantly enriched. REVIGO was used to summarize GO terms to reduce redundancy and group according to similarity (right labels).

By 12 h, there was an enrichment of downregulated transcripts associated with “positive regulation of intracellular signal transduction” in the gill which included several growth-related processes (Fig. 3b). By 24 h, the parent term “cytokine-mediated signaling pathway” was the largest enriched parent term and included the downregulation of transcripts associated with several GO terms related to cell growth and survival, including cell proliferation, apoptosis, and G2/M mitotic transition (Fig. 3b).

The bluegill gill’s response to TFM treatment involved expression changes of transcripts associated with immune function. For upregulated transcripts, “negative regulation of macrophage derived foam cell differentiation” (12 h) and “cellular response to transforming growth factor beta stimulus” (12 and 24 h) were the main responses (Fig. 3a). However, there was a large enrichment of GO terms associated with the immune response in downregulated transcripts in the gill (Fig. 3b). At 12 h, cytokine signaling, and T cell migration were enriched. By 24 h, the parent term “cytokine-mediated signaling pathway” included GO terms associated with immune function including “inflammatory response” and NF-ĸβ signaling (Fig. 3b). There was also an enrichment of downregulated transcripts associated with the parent terms “antigen presentation via MHC class I” and “viral life cycle” (Fig. 3b).

TFM exposure also affected metabolic processes in bluegill gills. For upregulated transcripts, this included enrichment of lipid metabolism (e.g., *abca1*, *cyp1a1*, *angptl4*; 12 h), gluconeogenesis (e.g., *pdk2*, *pgp*, *ptpn2*; 12 h), reactive oxygen species (ROS; e.g., *cdkn1a*, *slc25a33*, *thbs1*; 12 h), and glucose import (*insr*, *irs2*, *c1qtnf12*; 24 h). In downregulated transcripts, enriched metabolic processes included a regulation of primary metabolic processes and cellular amino acid metabolic processes (Fig. 3b). Tangentially, there was also an enrichment of GO terms in upregulated transcripts that were associated with a response to an organic compound (6h; Fig. 3a), which included the upregulation of several transcript targets, notably cytochrome P450s (*cyp1a1* and *cyp1b1*), caspases (*casp3* and *casp7*), and an aryl hydrocarbon receptor (*ahr*).

In the bluegill liver, TFM effects were generally restricted to transcripts pertaining to cellular growth, proliferation, and cellular death. Specifically, in upregulated transcripts, there was an enrichment of the parent GO terms “regulation of autophagy” (6 h), “positive regulation of apoptotic process” (12 h), “macroautophagy” (24 h), and “ubiquitin-dependent protein catabolism” (24 h; Fig. 4a). At 24 h, there was also enrichment of cell growth/death GO terms for downregulated transcripts including regulation of programmed cell death and cell cycle G2/M transitions (Fig. 4b). TFM also led to an enrichment of metabolic functions in the bluegill liver. Specifically, glycogen metabolic processes were enriched in downregulated transcripts which corresponded with lower differential expression of a glycogen debrancher enzyme (*agl*), α-glucosidase (*gaa*), glycogen synthase (*gys2*), and glycogen phosphorylase (*pygl*; Fig. 4b).

**Figure 4:**
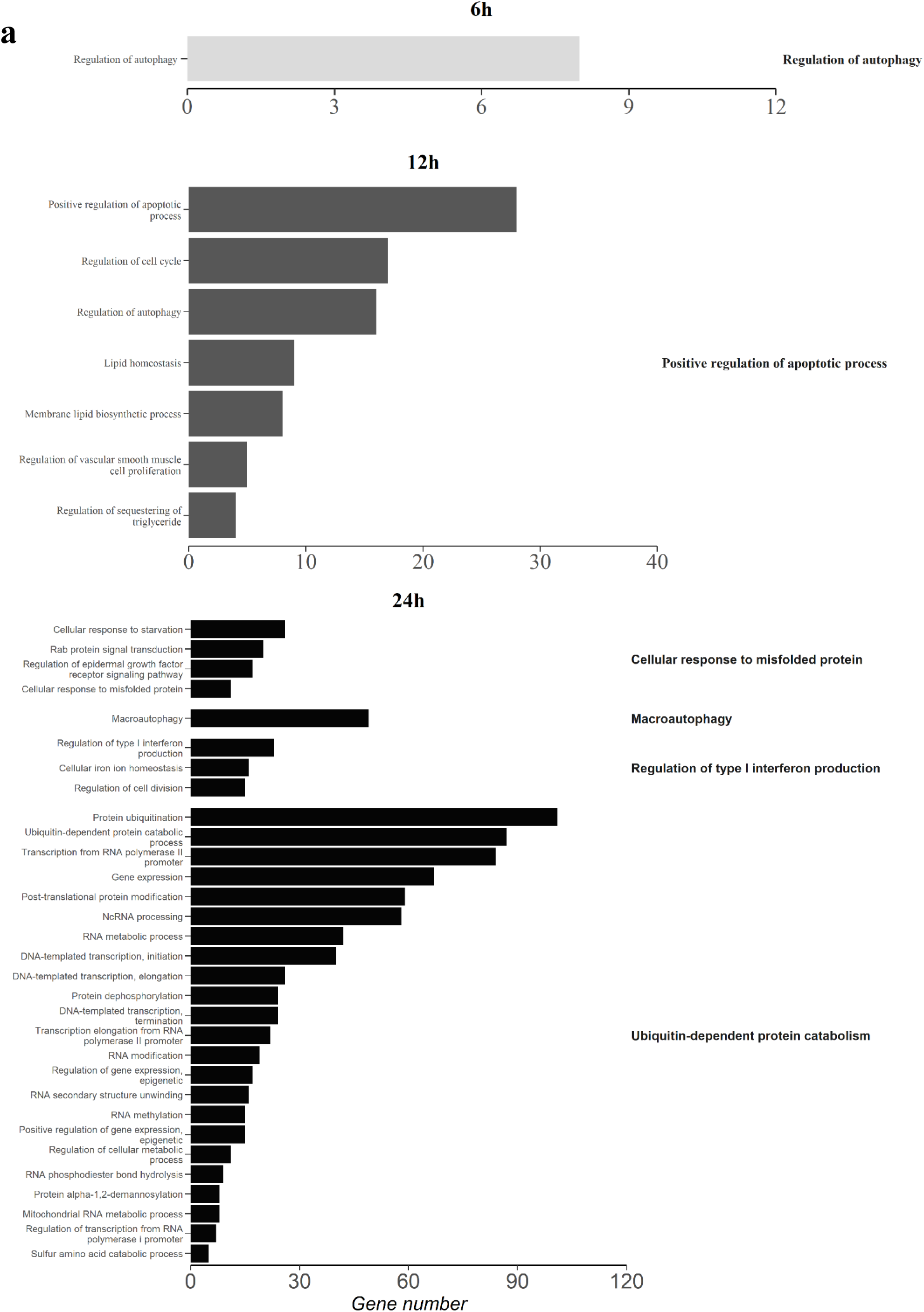
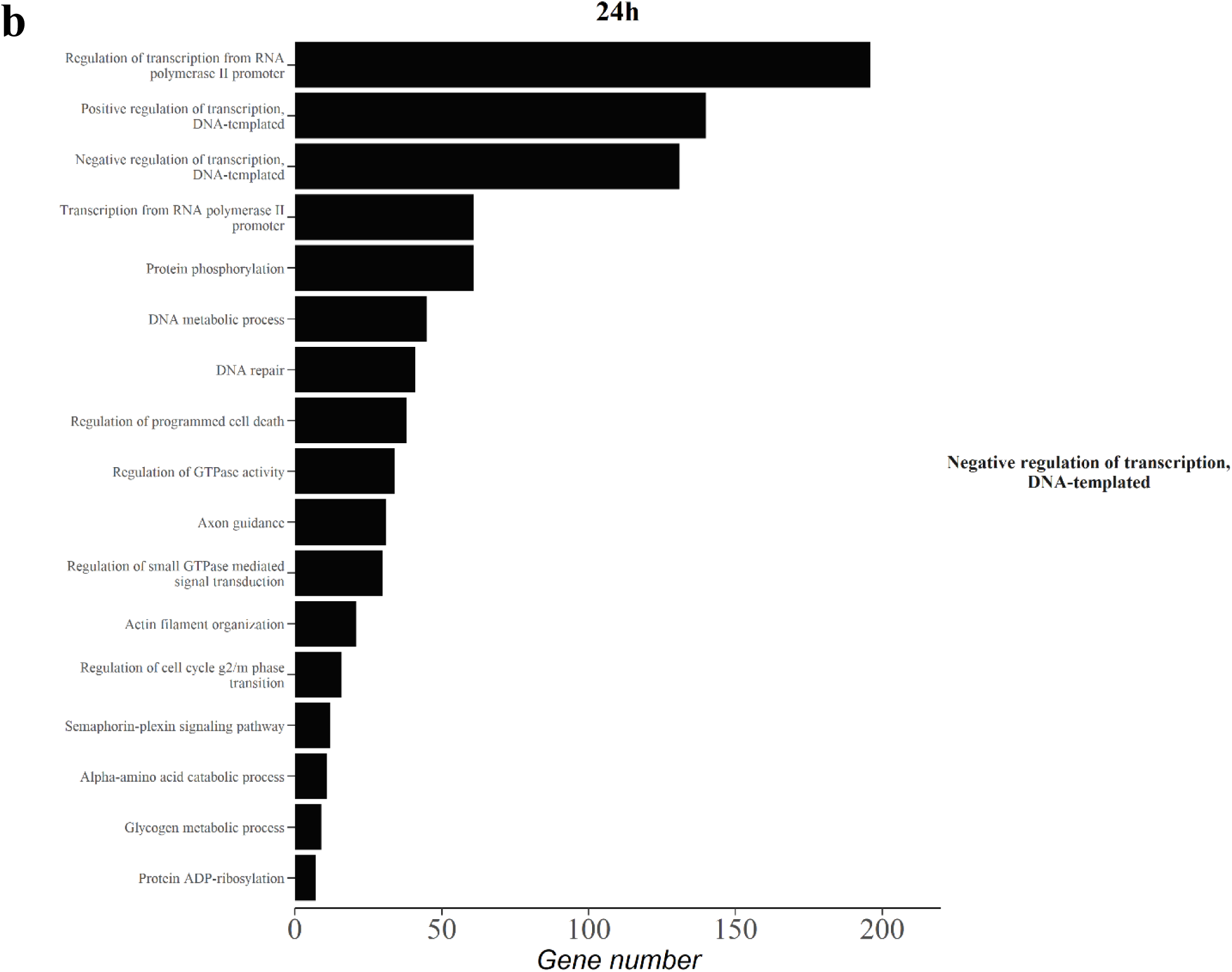
Summary of enriched gene ontology (GO) terms with transcripts that were upregulated (**a**) and downregulated (**b**) following 6 h (light grey), 12 h (dark grey) and 24 h (black) of TFM (3-trifluoromethyl-4’-nitrophenol; 22.06 mg L^-1^ nominal) exposure in the liver of bluegill (*Lepomis macrochirus*). Transcripts were considered differentially regulated at a false discovery rate < 0.05. Only GO terms from the functional analysis with an adjusted p < 0.05 and at least four transcripts were considered significantly enriched. REVIGO was used to summarize GO terms to reduce redundancy and group according to similarity (right labels).

## Discussion

Variation in the availability and functional status of different detoxification pathways plays a critical role in determining the interspecific and intraspecific sensitivity of animals to different pesticides^10^. Accordingly, we predicted that interspecific variation in the expression and diversity of transcripts coding for biotransformation enzymes would partially explain the differences in TFM tolerances between sea lamprey and bluegill. As predicted, we identified considerable interspecific variation in the transcriptomic responses to TFM exposure where all three suites of biotransformation transcripts (i.e., Phase I-III) were more diverse and responsive in bluegill relative to sea lamprey, suggesting a greater inherent ability to detoxify TFM. The differences in the transcriptome responses between the species supports an evolutionary basis of TFM tolerances in fishes. These results likely explain why >15-fold higher concentrations of TFM are required to produce a toxic effect in bluegill versus sea lamprey^3^, as well as the 10-fold higher 24 h LC10 observed in bluegill compared to sea lamprey^19^. A functional transcriptomics approach has therefore provided us with greater insight into the molecular basis for the different sensitivities of these two species to TFM, and how transcriptomic responses associated with detoxification genes explains these observations (Fig. 5).

**Figure 5:**
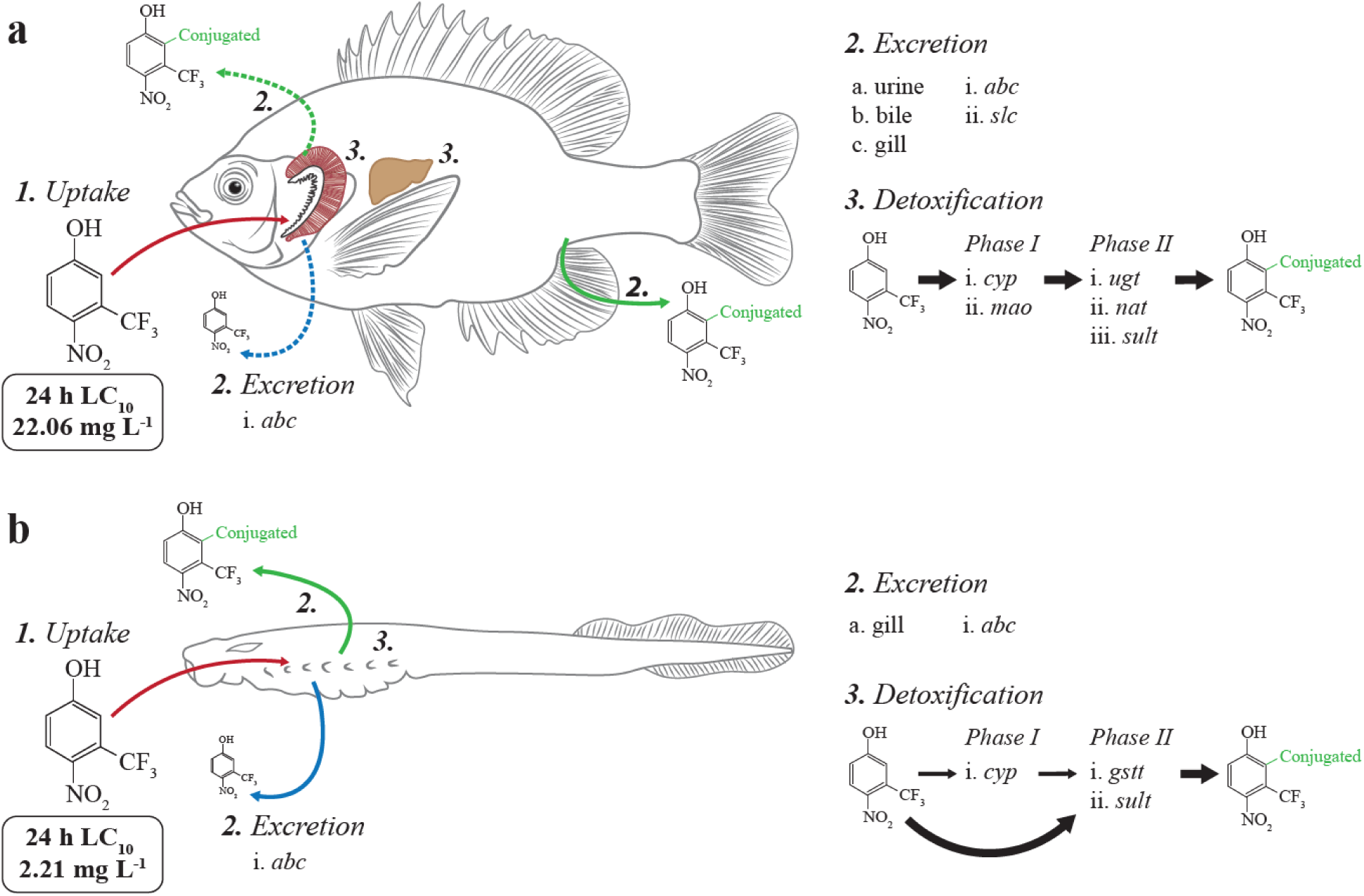
Overview of the pathways of TFM (3-trifluoromethyl-4’-nitrophenol) uptake, excretion, biotransformation, and elimination in bluegill (*Lepomis macrochirus*; **a**) and sea lamprey larvae (*Petromyzon marinus*; **b**). Detoxification transcripts that are presented here were found to have higher differential expression following exposure to TFM (bluegill = 22.06 mg L^-1^, lamprey = 2.21 mg L^-1^) over a 24 h period.

### Evolutionary considerations in lampricide tolerances

We found that differences in the expression and diversity of biotransformation transcripts could help to explain interspecific variation in TFM sensitivity among target sea lamprey and non-target teleost fishes. Specifically, differences in *ugt* transcript diversity and expression were likely a major factor dictating TFM tolerances in fishes. Variation in detoxification capacities have been exploited to improve the specificity of pesticides used across a broad range of agricultural, commercial, and domestic settings, while minimizing impacts on non-target species. For example, current agricultural practices make use of a combination of bioengineered crop plants that are tolerant to pesticides (e.g., Roundup Ready™ ^25^) as well as using taxon-specific pesticides (e.g., 2,4-dichlorophenoxyacetic acid in cereal crops^26^) to selectively control dicotyledons (broadleaf plants) without harming valuable monocotyledon crops. In contrast, many chemical insecticides (e.g., organophosphates and neonicotinoids) are broad-spectrum rather than selective pesticides, with detrimental non-target effects generally being reduced by applying them in a selective manner (e.g., via spot treatments or in baits)^27^. Thus, furthering our understanding of the genetic, physiological, and evolutionary mechanisms driving variation in pesticide sensitivities can allow for the development of more effective and targeted pesticide applications that minimize impacts on non-target species^2^. As pesticides used as lampricides cannot be applied in this manner, understanding genetic variation in detoxification mechanisms is key in protecting local biodiversity.

The sea lamprey control program in the Laurentian Great Lakes is perhaps the most notable and successful example of pesticides use in an invasive vertebrate^3^. High specificity was key in ensuring minimal impacts on non-target species as the pesticide was being administered directly into the natural environment. During the 1950’s, researchers conducted a large-scale series of toxicity tests, screening over 5,000 different compounds for an agent that could kill sea lamprey with higher selectivity than native species. Eventually, compounds containing a nitrophenol group were identified as the most selective agents, with TFM being the best substance^28^. Stream treatment with TFM has resulted in an estimated 90% reduction in sea lamprey populations in the Great Lakes^3^.

The exact mechanism underpinning the variation in TFM sensitivity among fishes has remained illusive. Phylogenetic variation in TFM sensitivity was apparent as closely related species (i.e., within order) demonstrated similarities in TFM toxicity values^3, 19^. Notably, native lampreys had comparable toxicity values (i.e., minimum lethal concentration 99% [MLC99]) to sea lamprey and were also adversely affected by lampricide treatments^29^, suggesting a taxonomic basis for the sea lamprey’s high sensitivity to TFM. Indeed, it is believed that Ugt’s are the principal enzyme in TFM detoxification as part of Phase II biotransformation ^3, 20, 21, 30^. In sea lamprey, Ugt activities and numbers of *ugt* gene isoforms appear to be low relative to teleost fishes^21–23^, suggesting an underlying genetic basis in dictating TFM tolerances. The sea lamprey’s phylogenetic position as a basal vertebrate could help explain the disparity in TFM tolerances. Lampreys diverged from the lineage leading to jawed vertebrates ∼500 million years ago, most likely after one round of whole genome duplication (WGD), with the second WGD occurring only in jawed vertebrates^31, 32^ and a third WGD in teleosts^31^; these WGD events are important in allowing the generation of new traits that can facilitate adaptation^31^. Consequently, this provided teleosts with a more diverse set of detoxification genes relative to lampreys, and perhaps even to non-teleost fishes like lake sturgeon, which are moderately sensitive to TFM (although less than lampreys^3^).

### Patterns of ugt expression

We show for the first time that there is a genetic basis underlying differences in TFM tolerances among target and non-target freshwater vertebrates. Specifically, bluegill had a larger diversity of *ugt’*s, compared to sea lamprey, and only bluegill showed differential expression of *ugt*’s in their gill and liver transcriptomes due to TFM exposure. In line with our predictions, we showed that differences in *ugt* expression patterns between bluegill and sea lamprey are likely one of the main factors driving interspecific variation in TFM tolerance in fishes.

Bluegill exhibited differential expression in the *ugt* gene family including *ugt1*, *ugt2*, and *ugt3*. In bluegill, *ugt3* appeared to be the most important *ugt* transcript in both tissues as it was expressed at the highest levels and underwent the greatest fold-increase (upwards of 28x), compared to *ugt1* and *ugt2*. *ugt3* genes have not been identified in fishes previously and was only recently discovered in humans with their functional role remaining uncertain^33^. However, the Ugt3 enzyme appears to be involved in detoxifying polyaromatic hydrocarbons, and 4- nitrophenol in mammals^33^, suggesting a role in TFM detoxification in bluegill as well. We also noted an upregulation of *ugt2b9* in the bluegill liver. Thus, hepatic biotransformation using *ugt3* and, to a lesser extent, *ugt2b9* may be a key factor in the high tolerance of bluegill to TFM compared to sea lamprey and perhaps other teleost fishes^3^.

### Insight into detoxification pathways using transcriptomics

Variation in the detoxification transcriptome between bluegill and sea lamprey was evident in genes associated with Phase I biotransformation, predominantly genes coding for *cyp*’s, which can detoxify organic compounds^24^. These enzymes may have a role in TFM detoxification as evident by the presence of Cyp-conjugated TFM metabolites in fishes^21^, but their relative importance compared to Phase II processes is uncertain. In both bluegill and sea lamprey, *cyp1a1* and *cyp1b1* appeared to be the dominant *cyp* transcripts that responded to TFM, which is unsurprising given that *cyp1* genes are involved in the detoxification of a diversity of organic compounds^24^. Interestingly, bluegill had a far greater magnitude of response, as evident by the high fold-changes in expression (upwards of ∼187x), and a greater diversity of differentially expressed *cyp* transcripts, compared to sea lamprey. However, it is unclear how important Cyp’s are for detoxifying TFM, as no studies have detected the corresponding Phase I metabolites *in vivo*, as compared to the Phase II sulfate and glucuronide conjugates^21, 22, 34^.

Further studies, both *in vitro* and *in vivo*, are required to better understand the functional role of Cyp’s in TFM biotransformation in both species.

Phase II biotransformation transcripts also showed a clear taxonomic divergence that may help explain interspecific variation in TFM tolerances. Bluegill exhibited an upregulation of sulfotransferases (N=3) and N-acetyltransferases (N =2) across both tissue types, whereas only a single sulfotransferase and glutathione S-transferase responded in sea lamprey gill. As sulfotransferases are important in TFM detoxification^3, 21^, the greater capacity to conjugate via sulfation as well as through glucuronidation in bluegill likely supports high TFM metabolism when compared to sea lamprey. The presence of N-acetyltransferases in bluegill also suggests that TFM is undergoing acetylation^24^, which has been characterized in bluegill previously^21^. Our results suggest a taxonomic disparity in Phase II biotransformation pathways involved in TFM detoxification, with bluegill able to upregulate several Phase II biotransformation processes to effectively eliminate TFM compared with sea lamprey.

Interspecific variation was also evident in Phase III biotransformation processes, which are mainly involved in the transport and elimination of toxicants^24^. We detected a divergent effect of TFM exposure on Phase III transcripts where numbers of responding *abc* and *slc* transcripts were greater in sea lamprey and bluegill, respectively. This might suggest a disparity in TFM elimination routes given that Abc’s are generally active transporters while Slc’s are bidirectional passive transporters^24^. In the lamprey, the high *abc* response, as well as the general lack of transporter expression in the liver, may suggest a predominantly branchial-mediated elimination^3^ as any TFM excretion would be against a concentration gradient (i.e. high environmental TFM concentration). In bluegill, the high environmental concentrations of TFM (∼21 mg L^-1^ [101 µM] in water, 23 nmol g^-1^ ww in liver^19^) coupled with the lower branchial *abc* expression, relative to lamprey, suggests that TFM elimination can be through biliary/renal excretion as previously reported in teleosts^3, 30^. Together, this appears to suggest that bluegill have a more responsive and effective system for TFM elimination.

### GO term enrichment and tissue responses to TFM

In fishes, studies of TFM toxicity have largely been restricted to characterizations of energy metabolism and mortality^17, 19^. However, we found several biomarkers associated with inhibited cell cycle progression and growth, and higher levels of ubiquitination and apoptosis, in both bluegill and sea lamprey, suggesting that TFM’s effects extend beyond simply uncoupling oxidative phosphorylation^16, 35^, indicating a new mode of TFM toxicity in fishes. Increases in ROS or oxidative damage are likely the primary mechanism by which mitochondrial uncouplers exert tissue-level repression of growth^36^. While TFM effects on ROS generation have not been quantified, TFM does increase mitochondrial respiration rates^16^, which may serve to elevate cellular ROS levels^37^. Coupled with the GO term enrichment patterns, our results suggest that TFM-induced ROS is likely mediating cellular arrest as seen in niclosamide^38, 39^ and 2,4- dinitrophenol (DNP, a TFM analogue)^40, 41^ exposures. While fish-specific examples are limited, zebrafish (*Danio rerio*) exposed to 4-nitrophenol experienced reduced cellular growth and proliferation, and heightened cell death^36^, aligning with our results. At a broader scale, increases in oxidative damage by ROS can impair cellular growth and cell cycle progression as well as inducing apoptosis in fishes^42, 43^. Together, our results suggest that TFM has a potential role in generating oxidative damage in the cell.

Despite TFM toxicity primarily affecting energy metabolism^3^, enrichment of these metabolic GO terms was limited. In sea lamprey, there was a negative regulation of GO terms associated with lipid storage, gluconeogenesis, and insulin responsiveness. Likely, these changes were needed to increase energy mobilization for supporting heightened metabolic demands under TFM exposure^17–19^. In bluegill, metabolic changes included positive regulation on gluconeogenesis, glucose import, and lipid metabolism and negative regulation of glycogen metabolism suggesting that increased energetic expenditure during TFM exposure is likely occurring, probably due to increased rates of mitochondrial respiration and detoxification costs^19^. Also, as predicted, the effects of TFM on both species’ transcriptomes was more apparent with longer exposure durations (i.e., more differentially expressed transcripts) suggesting an exhaustion of detoxification systems.

## Conclusions

Here, we showed how transcriptome responses demonstrate interspecific variation in pesticide sensitivities, which are critical in developing more targeted invasive species control. We identified differences in *ugt* diversity and responsiveness to TFM, which appeared to be linked to differences in TFM sensitivities between bluegill and sea lamprey highlighting the importance of evolutionary history in mediating toxicity resilience. In the context of sea lamprey control, this knowledge will be useful in developing a predictive framework for assessing community-scale impacts of lampricides. More broadly, similar principles are likely to apply to a wide range of systems where invasive species control efforts use pesticides extensively. By identifying key genes involved in detoxification in native fauna, we can develop methods for addressing the relative impacts that pesticide treatments have on a system. For TFM specifically, this would likely include developing screenings of Ugt’s, Cyp’s, and Sult’s as potential biomarkers of exposure. This approach could help shed light on more subtle interspecific differences in sensitivity to TFM even among teleosts. As invasive species control efforts seek to provide more effective and targeted pesticide applications^13^, furthering our understanding of the toxicological mode of action and evolutionary variation in tolerances among species is likely to greatly improve the use of pesticides.

## Methods

### Animal care and holding

Fish collection and holding conditions are presented in Lawrence et al.^19^. Briefly, juvenile bluegill (*n* = 200; total length [TL] = 97.3 ± 0.88 mm; mass = 25.51 ± 0.75 g) were sourced from Kinmount Fish Farm (Kinmount, ON, Canada) in September 2018. Bluegill were transported to the animal holding facility at Wilfrid Laurier University (Waterloo, ON, Canada; ∼1000 L holding tank; ∼ 0.5 L min^-1^; T = 12–14°C; pH 8.1–8.2, alkalinity ∼255 mg L^-1^ as CaCO_3_), where they were provided with a combination of commercial fish feed (EWOS #1, Cargill, ON, Canada) and bloodworms daily.

Larval sea lamprey (*n* = 568; TL = 104.94 ± 0.69 mm; mass = 1.47 ± 0.03 g) were electrofished from a tributary of Lake Huron by the United States Fish and Wildlife Service in April 2018 and were temporarily stored at the Hammond Bay Biological Station (Millersburg, MI, USA). Sea lamprey were then transported to Wilfrid Laurier University and held under similar water conditions to the bluegill, although the lamprey tank was smaller (∼100 L) and contained a 8–10 cm deep layer of sand to facilitate natural burrowing^44^. Lamprey were fed on a diet of baker’s yeast weekly (see Lawrence et al. ^19^ for specific details). All experimental series were conducted under approval from the Wilfrid Laurier University Animal Care Committee (Animal Use Protocol No. R18001) under the guidelines established by the Canadian Council of Animal Care.

### TFM exposures

Larval sea lamprey and bluegill were exposed to either control conditions (i.e., no toxicant) or field grade TFM (35% active ingredient dissolved in isopropanol; Clariant, Griesheim, Germany) for up to 24 h. The TFM exposure concentrations were equivalent to the species-specific concentration of TFM that was lethal to 10% of each species over 24 h (24-h LC_10_), previously determined for the cohorts of sea lamprey (24-h LC_10_ = 2.21 mg L^-1^) and bluegill (24-h LC_10_ = 22.06 mg L^-1^)^19^. Exposures were performed in triplicate with animals being held in glass aquaria (10 L for sea lamprey larvae, 14 L for bluegill) in groups of 2–3 or 6 individuals per tank for bluegill and sea lamprey larvae, respectively. The TFM exposures were static with temperature being maintained at the fish’s acclimation temperature (∼14–15°C).

Tissue sampling at 6 h, 12 h, and 24 h of exposure. At these discrete intervals, fish were netted and placed one at a time into buffered tricaine methanesulfonate (MS-222; Syndel, Nanaimo, BC, Canada; 1.5 g L^-1^ MS-222 with 3.0 g L^-1^ NaHCO_3_) to euthanize the animals. The livers and gill of both species were excised, and placed into RNA*later* (Invitrogen, ThermoFisher Scientific, Mississauga, ON, Canada), held at 4°C for at least 24 h, and then stored at -80°C for later RNA extraction.

### RNA extraction and transcriptome sequencing

Extractions of total RNA were performed using a commercially available kit (RNeasy Plus Mini Kit; Qiagen, Toronto, ON, Canada) according to the manufacturer’s specifications. Following extraction, an initial quality control check was performed using the 260/280 absorbance ratio and the 260/230 absorbance ratio with a NanoDrop One Microvolume UV-Vis Spectrophotometer (ThermoFisher, Mississauga, ON, Canada). Quality was further assessed using a Qubit RNA IQ assay (ThermoFisher, Mississauga, ON, Canada), read on a Qubit 4 fluorometer (ThermoFisher, Mississauga, ON, Canada), as a final check of RNA integrity. Samples were then diluted to a standardized volume (50 ng µL^-1^) and stored at -80°C until shipment to the sequencing facility.

RNA-seq library preparation and sequencing were performed by the Centre d’Expertise et de Services Génome Québec (Montreal, QC, Canada). Before sequencing, samples were again verified for integrity using both spectrophotometry (NanoDrop ND 1000; ThermoFisher, Mississauga, ON, Canada) and electrophoresis (Bioanalyzer 2100; Agilent, Santa Clara, CA, USA; RIN ranges: sea lamprey, 8.4–10.0; bluegill 8.5–9.8). Sequencing of cDNA libraries was achieved using the NovaSeq 6000 sequencing system (Illumina, Vancouver, BC, Canada). Total sequencing read counts can be found in Table S1.

### Transcriptome assembly, alignment, and annotation

#### RNA-seq QC

Raw reads were downloaded from Genome Quebec for processing and downstream analysis on a University of Manitoba personal Linux Server, as well as supercomputers belonging to Westgrid (Grex) and Compute Canada (Beluga, Cedar, and Graham). Fastqc (v 0.11.9) was run on the raw sequencing data to assess quality, and MultiQC (https://github.com/ewels/MultiQC) was used to generate reports from the Fastqc for easier viewing across samples^45, 46^. It is important to note here that RNA-seq data files contained sequencing data from fish exposed to TFM as well as those exposed to niclosamide or a mixture of TFM and niclosamide. The latter two exposures are part of a complementary but ongoing study and are not included in the final results here. Raw sequencing quality and duplication rates met our expectations across all 148 sea lamprey samples and all 163 bluegill samples (combined numbers of gill and liver tissue for each species that were obtained from the exposure series). Adapters were trimmed from the raw sequences using Trimmomatic (v0.36), Fastqc and MultiQC were run again to ensure successful trimming^47^.

#### *De novo* transcriptome assembly

Since bluegill does not have a reference genome to map reads back to, *de novo* transcriptome assembly was carried out for both bluegill and sea lamprey to keep methods consistent^48^. These *de novo* assemblies were each constructed from a list containing a single library for each tissue and timepoint. These input libraries were selected based on the highest read count, accounting for duplication rates for each condition. For bluegill (*n* = 26), we used one paired-end library from liver and gill for each of the following treatments and timepoints: 6, 12, and 24 h for niclosamide (*n* = 6), TFM (*n* = 6) and TFM:niclosamide mixture (*n* = 6), as well as 0, 6, 12, and 24 h for the control (*n* = 8). For lamprey (*n* = 22), paired-end libraries from liver and gill for 6, 12, and 24 h for niclosamide (*n* = 6), TFM (*n* = 6), and control (*n* = 6) were used, but the TFM:niclosamide mixture only had libraries at 6 and 12 h (*n* = 4), since the combination killed 93% of all lamprey by 12 h of exposure (see Lawrence et al.^19^).

Each assembly was generated using Trinity (v2.8.5), run with the following command: Trinity --max_memory 250G --seqType fq --samples_file samples.txt --KMER_SIZE 25 -- SS_lib_type RF --CPU 24 --bflyCalculateCPU --output full_trinity_assembly &>log_RF_2.txt &^49^. Following Trinity, the completeness of the assemblies was assessed using BUSCO (v4.1.4) to test for the presence of Benchmarking Universal Single Copy Orthologs that are conserved within all eukaryotes using the eukaryota_odb10 database^50^. The bluegill assembly was 98.4% complete with one fragmented and three missing BUSCOs out of the database of 255. The sea lamprey was 96.1% complete with six fragmented and four missing BUSCOs (Table 1). These results are comparable to a recently published *de novo* larval sea lamprey transcriptome assembled from muscle, liver, and brain. The bluegill assembly had 559,416,397 assembled bases in 498,997 transcripts with a median contig length of 413, contig N50 of 2,992, and %GC of 44.38. The sea lamprey assembly had 377,365,173 assembled bases in 474,000 transcripts with a median contig length of 358, contig N50 of 1,648, and %GC of 53.10.

#### *De novo* transcriptome annotation

Trinity assemblies were processed with the Trinotate (v3.2) annotation protocol (http://trinotate.github.io), which incorporates many bioinformatics tools, namely TransDecoder (v5.5.0), BLASTP (v 2.10.0), BLASTX (v2.10.0), HMMER (packaged in Trinotate v3.2), TMHMM (v2.0c), SignalP (v4.1), and RNAmmer (packaged in Trinotate v3.2) to identify open reading frames (ORFs) within the assembly and test these ORFs against databases of known proteins, mRNA transcripts, transmembrane helices, signal peptides, and ribosomal RNA sequences in order to provide candidate annotations and gene ontology (GO) terms for transcripts within the assembly^49, 51–55^. Predicted annotations were used in downstream differential expression analyses.

#### Super transcriptome assembly

Although *de novo* assemblies are often essential when reference genomes do not exist, carrying out differential expression analyses on raw assemblies is complicated by the fact that multiple transcripts are often generated for each gene. To combat this issue, we used the Corset- Lace pipeline (https://github.com/Oshlack/Lace) to generate SuperTranscripts from our *de novo* assemblies. First, we mapped reads back to the Trinity assembly using Bowtie2, then carried out hierarchical clustering of transcripts based on shared reads and expression across transcripts, finally creating an assembly of SuperTranscripts based on clustering of gene groups^56–58^. For bluegill, Corset-Lace reduced 498,997 transcripts to 109,702 SuperTranscripts, and 474,000 transcripts were reduced to 129,219 SuperTranscripts in sea lamprey. These SuperTranscriptomes were run through BUSCO, as above, and bluegill SuperTranscriptome displayed 84.7% completeness, whereas the lamprey reported 91% completeness. Although both of these measures are slight decreases from the original 98.4% and 96.1% for bluegill and lamprey, respectively, the biggest change in the BUSCO report was the number of complete and single-copy BUSCOs compared to the number of complete and duplicated BUSCOs (Table S2).

Given that the genes in the BUSCO list represent Benchmarking Universal Single-Copy Orthologs across eukaryotes, the high percentage of “complete duplicated BUSCOs (D)”, 58.8% and 60.4% in bluegill and lamprey, respectively, indicated that our *de novo* Trinity assemblies contained numerous transcripts for the same gene, which would be problematic when assessing differential expression. Our SuperTranscriptomes decreased the overall transcript number in our assemblies by ∼3.4–4.5 times, slightly decreased the overall percentage of complete BUSCOs (the sum of single-copy and duplicated BUSCOs) by ∼1.05–1.15 times, but most importantly, decreased the number of duplicated BUSCOs by ∼17–21 times. The Corset-Lace pipeline greatly increased the number of complete single-copy BUSCOs (S) for both species. Although the Corset-Lace pipeline resulted in slightly fewer genes overall, we believe that those remaining were better assembled, with far fewer genes present across multiple transcripts in the assembly, which increased our power for differential expression analyses.

### Differential expression analyses

Reads from bluegill and lamprey TFM experimental and control conditions were mapped to the corresponding SuperTranscriptome using STAR (v2.6.1a)^59^. The resulting BAM files from STAR were used to generate transcript counts using featureCounts (part of subread v2.0.0)^60^ for both exon and transcript-based analyses.

Transcriptomic analysis was performed in R studio (v1.3.1093) using the R programming language (v3.5.1)^61^. Principal analyses of differential expression patterns were performed using the package ‘edgeR’ (v3.24.3)^62, 63^ using a quasi-likelihood pipeline^64^. Count data were first filtered to remove lowly expressed genes with the “filterByExpr” function which used the experimental design matrix to determine minimum gene count thresholds. Filtered count data were then normalized using the trimmed mean of M-values^65^. At this point, normalized data were visually inspected using principal component analysis (PCA; packages ‘FactoMineR’^66^ and ‘factoextra’^67^; see Fig. S1) and a multi-dimensional scaling (MDS) plot (package ‘Glimma’^68^). Individual samples were also visually inspected using a mean difference (MD) plot.

To determine patterns of differential expression, count data were first modeled against a negative binomial distribution^63^. From this, we obtained dispersal estimates using a Cox-Reid profile adjusted likelihood method^63^, which were visually inspected by use of a biological coefficient of variation (BCV) plot. Quasi-likelihood (QL) dispersion estimates were determined and were subject to a QL F-test to determine transcripts that had statistically significant differential gene expression^69^. From the resulting QL F-tests, we pulled out pairwise contrasts of interest which include control versus TFM exposed fish for each timepoint of exposure (i.e., 6, 12, 24 h). Statistical significance of pairwise comparison was accepted at α = 0.05 where all *p-* values were adjusted for false discovery rates (FDR) via a Benjamini-Hochberg correction^70^.

### Enrichment analyses

One of our primary goals was to ascertain specific pathways and processes that were underlying the transcriptome response to TFM exposure. To do so, we conducted analyses in pathway enrichment using the R package ‘enrichR’ (v2.1)^71^, which allows R access to the Enrichr databases (https://maayanlab.cloud/Enrichr/)^72^. This platform compares differentially expressed genes in a dataset and determines a list of GO terms that are enriched under the experimental treatment^72^. Here, we used three main databases for comparison: GO_Biological_Process_2018, GO_Molecular_Function_2018, and GO_Cellular_Component_2018. Before making database comparisons, unannotated superTranscript clusters were removed from the analysis. For annotated transcripts, UniProt gene names were retrieved using the UniProt’s Retrieve/ID mapping tool (https://www.uniprot.org/uploadlists/). In each transcripts list enrichR was run and GO term lists were subsequently filtered for those that were statistically significant (adjusted *p*-value < 0.05) and had four or more genes associated with a particular GO term^73^.

To identify general patterns and biological features of interest, we simplified the resulting enrichR GO term lists using a web-based platform, REVIGO (http://revigo.irb.hr/)^74^, which summarizes GO term patterns by grouping terms based on their similarity and then creates a hierarchical structure of the terms^74^. Our REVIGO analyses were at a similarity level of 0.5 (i.e., small) and used the adjusted *p*-values generated from enrichR. Treemap files from the REVIGO output were used to generate GO term enrichment plots in ggplot2^75^. To reduce term redundancy, we opted to only use biological processes and molecular functions from the REVIGO analysis. In cases where the biological processes and molecular functions provided functionally similar results, only the biological processes were displayed in the Results section with the molecular functions being available in the Supplementary Materials. Similarly, summary plots that offered no insight on specific processes (e.g., DNA replication, protein kinase activity, GTPase activities, etc.) were placed into Supplementary Materials.

### Identification of differentially expressed detoxification transcripts

We next sought to identify specific mechanisms underlying TFM detoxification and understand species-specific differences in detoxification ability (see Lawrence et al.^19^ for a review). To achieve these goals, we also incorporated a targeted approach, which required us to identify known and suspected candidates for TFM detoxification, including *ugt*’s, one of the principal enzymes believed to be involved in Phase II TFM detoxification (see Wilkie et al.^3^), as well as other enzymes/proteins involved with Phases I, II, and III of detoxification. In the case of *ugt* transcripts, *ugt*-associated transcripts were first identified in each species’ annotated transcriptome. These transcripts were then used to filter each list of species- and tissue-specific differentially expressed transcripts to examine how these enzymes responded to TFM exposure.

In the case of the Phase I–III proteins, we opted to filter a broad list of common protein families involved in detoxification from the differentially expressed transcripts lists and then manually check their functional role based on their UniProt IDs. As discussed in the Results, we filtered transcripts in the transcriptome associated with detoxification, xenobiotic removal, and/or organic compound breakdown. In the case of Phase III transcripts, we also included transporters that were involved in organic ion transport and those transporting UDP/glucuronide substrates. Phase I search terms included cytochromes P450 (CYPs), alcohol dehydrogenase (ADH), aldehyde dehydrogenase (ALDH), monoamine oxidases (MAO), and paraoxonases (PON). Phase II protein search terms included sulfotransferases (SULT), glutathione transferases (GST), glycine N-methyltransferase (GLYAT), N-terminal acetyltransferases (NAT), and methyltransferases (MT). Phase III search terms included solute carrier’s (SLC) and ATP- binding cassette (ABC) transporters.

#### Code and Data Availability

All commands and scripts used for quality control (QC), assembly, annotation, read mapping, transcript counting and RNAseq analyses are available at https://github.com/phil-grayson/transcriptome_lawrence_jeffries/. The raw sequence reads are available at the National Center for Biotechnology Information Sequence Read Archive (accession number SUB8714632). Gene lists and GO term list can be found in the Supplementary Materials.

## Acknowledgements

Funding for the research was provided by the Great Lakes Fishery Commission (2018_JEF_54072). The authors would like to thank Dr. Oana Birceanu for logistical assistance, Fisheries and Oceans Canada for providing the TFM used in this study, Darren Foubister and Josh Sutherby for conducting the exposures, and Drs. Sara Good and Matthew Doering, who provided helpful discussion and comments for completing the *de novo* assembly and the gene annotations, and the crew at the U.S. Geological Survey’s Hammond Bay Biological Station for collecting the larval lamprey used in this study.

## Contributions

KMJ, MPW, RGM, JMW, CJG, and MFD were responsible for the experimental design. The labs of KMJ and MPW carried out the exposures and sample collection. Data and sample analyses were conducted by MJL, PG, and JDJ. The first draft of the manuscript was written by MJL and PG, with all authors contributing to the refinement and production of the final manuscript.

---

**Figure S1:**
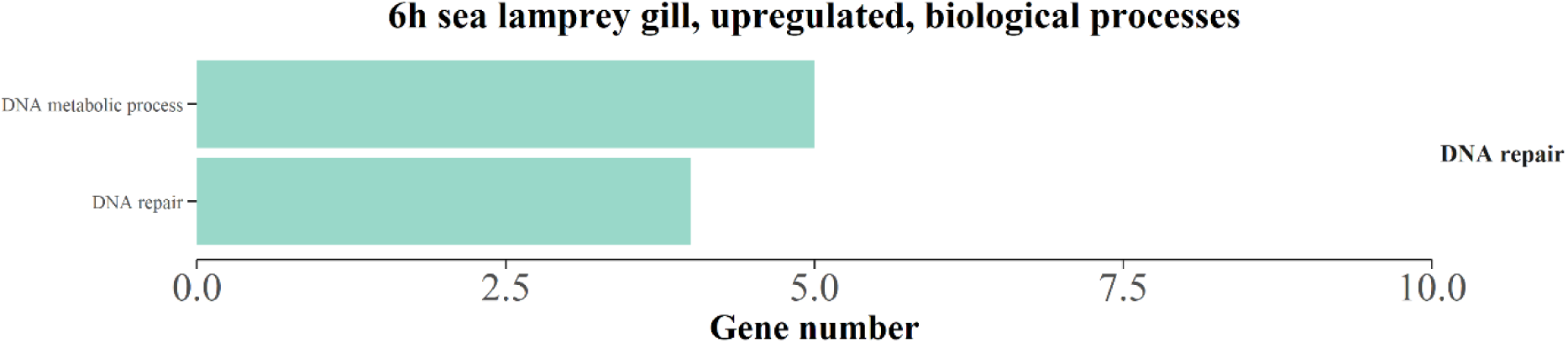
Summary of enriched gene ontology (GO) terms associated with biological processes in transcripts that were upregulated following 6 h of TFM (3-trifluoromethyl-4’-nitrophenol; 2.21 mg L^-1^ nominal) exposure in the gills of sea lamprey (*Petromyzon marinus*) larvae. Transcripts were considered differentially regulated at a false discovery rate < 0.05. Only GO terms from the functional analysis with an adjusted p < 0.05 and at least four transcripts were considered significantly enriched. REVIGO was used to summarize GO terms to reduce redundancy and group according to similarity (right labels).

**Figure S2:**
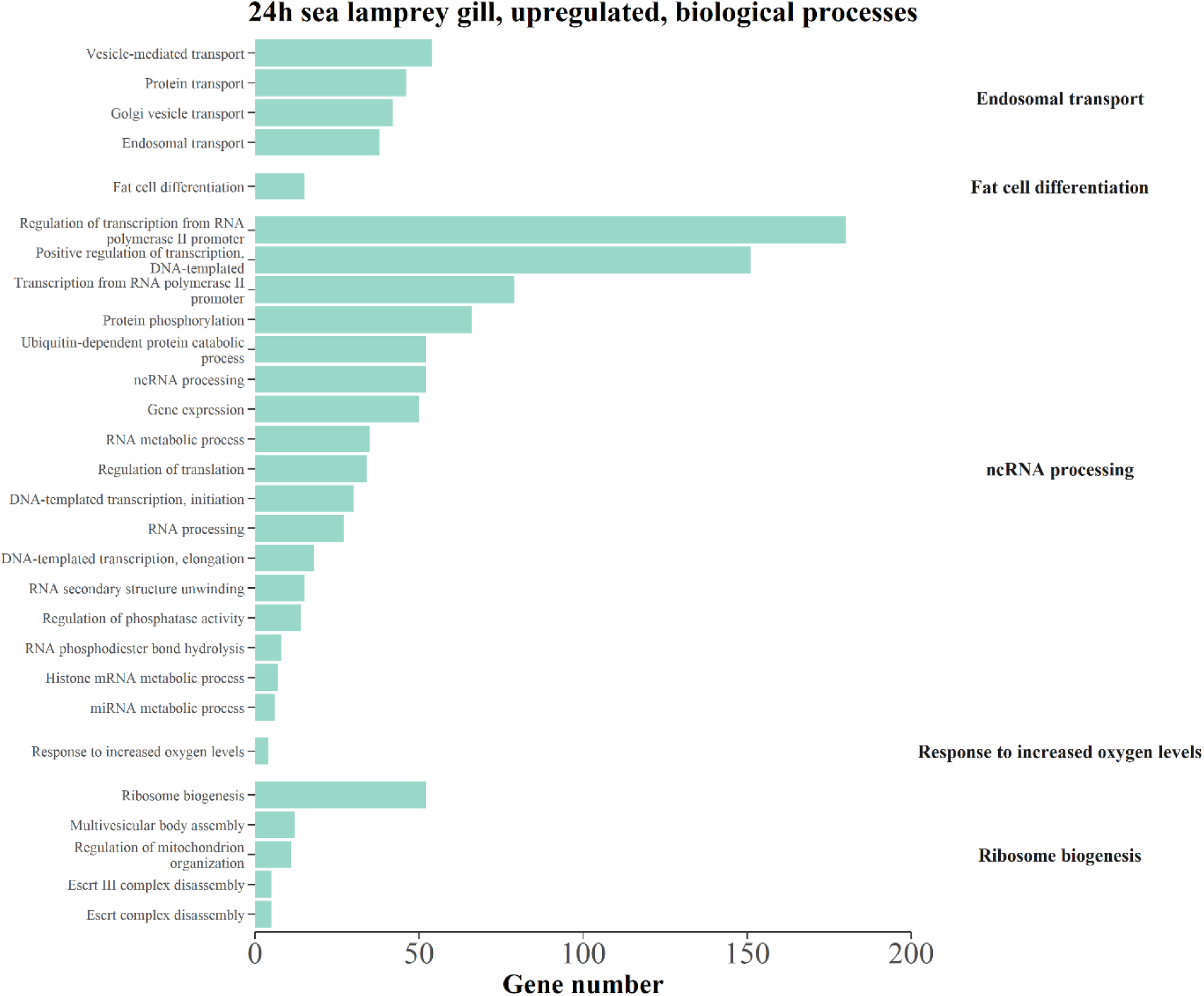
Summary of enriched gene ontology (GO) terms associated with biological processes in transcripts that were upregulated following 24 h of TFM (3-trifluoromethyl-4’-nitrophenol; 2.21 mg L^-1^ nominal) exposure in the gills of sea lamprey (*Petromyzon marinus*) larvae. Transcripts were considered differentially regulated at a false discovery rate < 0.05. Only GO terms from the functional analysis with an adjusted p < 0.05 and at least four transcripts were considered significantly enriched. REVIGO was used to summarize GO terms to reduce redundancy and group according to similarity (right labels).

**Figure S3:**
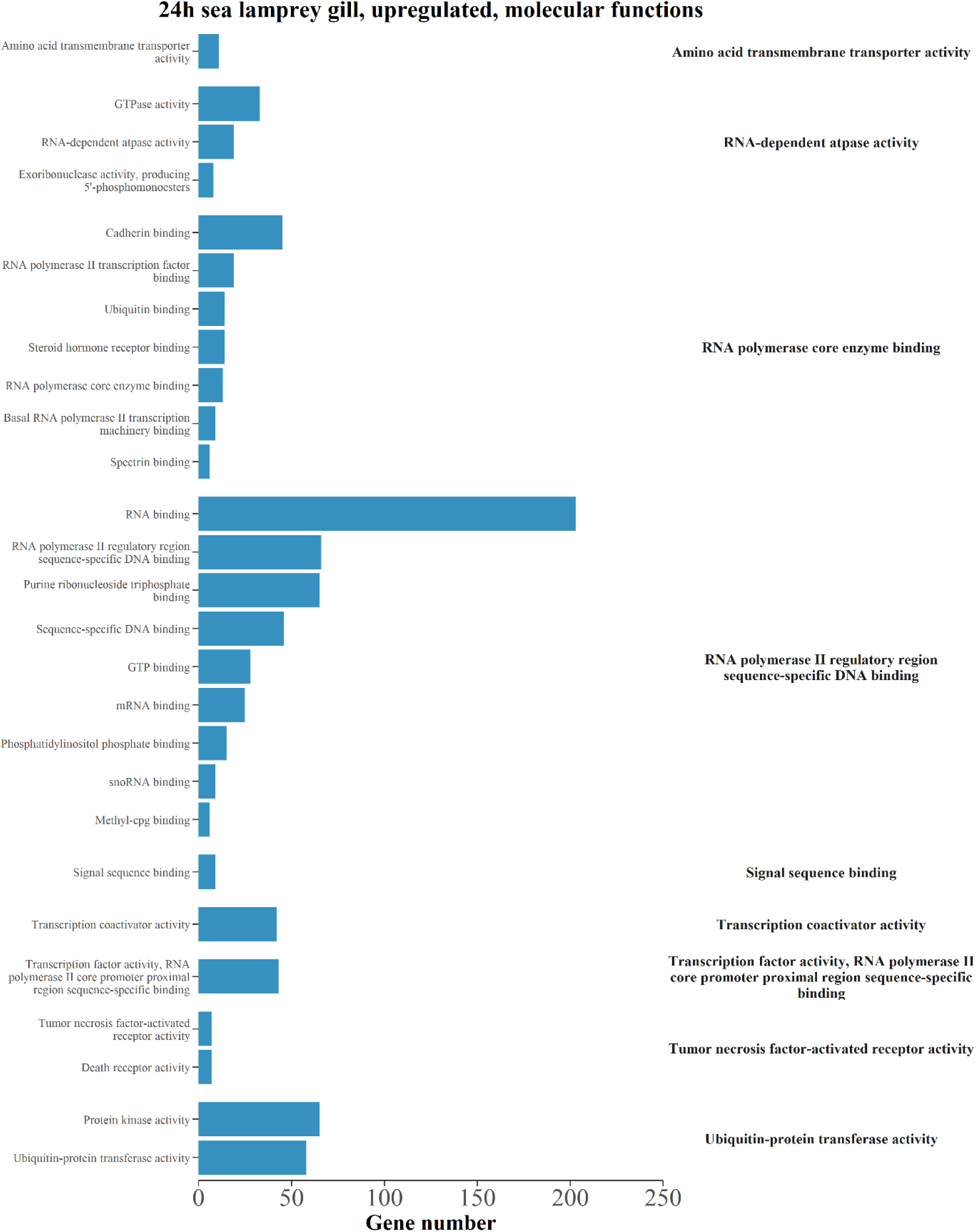
Summary of enriched gene ontology (GO) terms associated with molecular functions in transcripts that were upregulated following 24 h of TFM (3-trifluoromethyl-4’-nitrophenol; 2.21 mg L^-1^ nominal) exposure in the gills of sea lamprey (*Petromyzon marinus*) larvae. Transcripts were considered differentially regulated at a false discovery rate < 0.05. Only GO terms from the functional analysis with an adjusted p < 0.05 and at least four transcripts were considered significantly enriched. REVIGO was used to summarize GO terms to reduce redundancy and group according to similarity (right labels).

**Figure S4:**
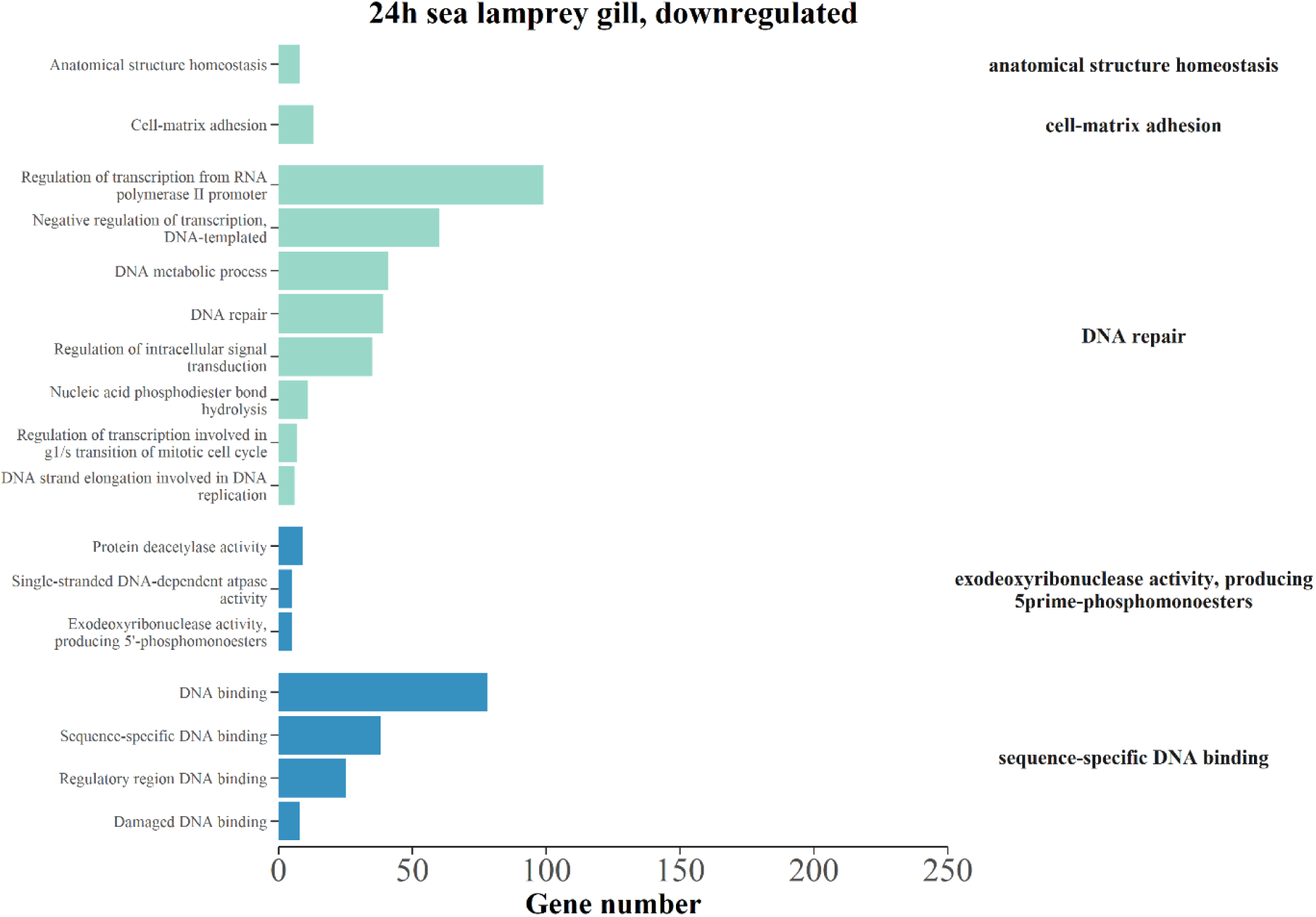
Summary of enriched gene ontology (GO) terms associated with biological processes (green) and molecular functions (blue) in transcripts that were downregulated following 24 h of TFM (3-trifluoromethyl-4’-nitrophenol; 2.21 mg L^-1^ nominal) exposure in the gills of sea lamprey (*Petromyzon marinus*) larvae. Transcripts were considered differentially regulated at a false discovery rate < 0.05. Only GO terms from the functional analysis with an adjusted p < 0.05 and at least four transcripts were considered significantly enriched. REVIGO was used to summarize GO terms to reduce redundancy and group according to similarity (right labels).

**Figure S5:**
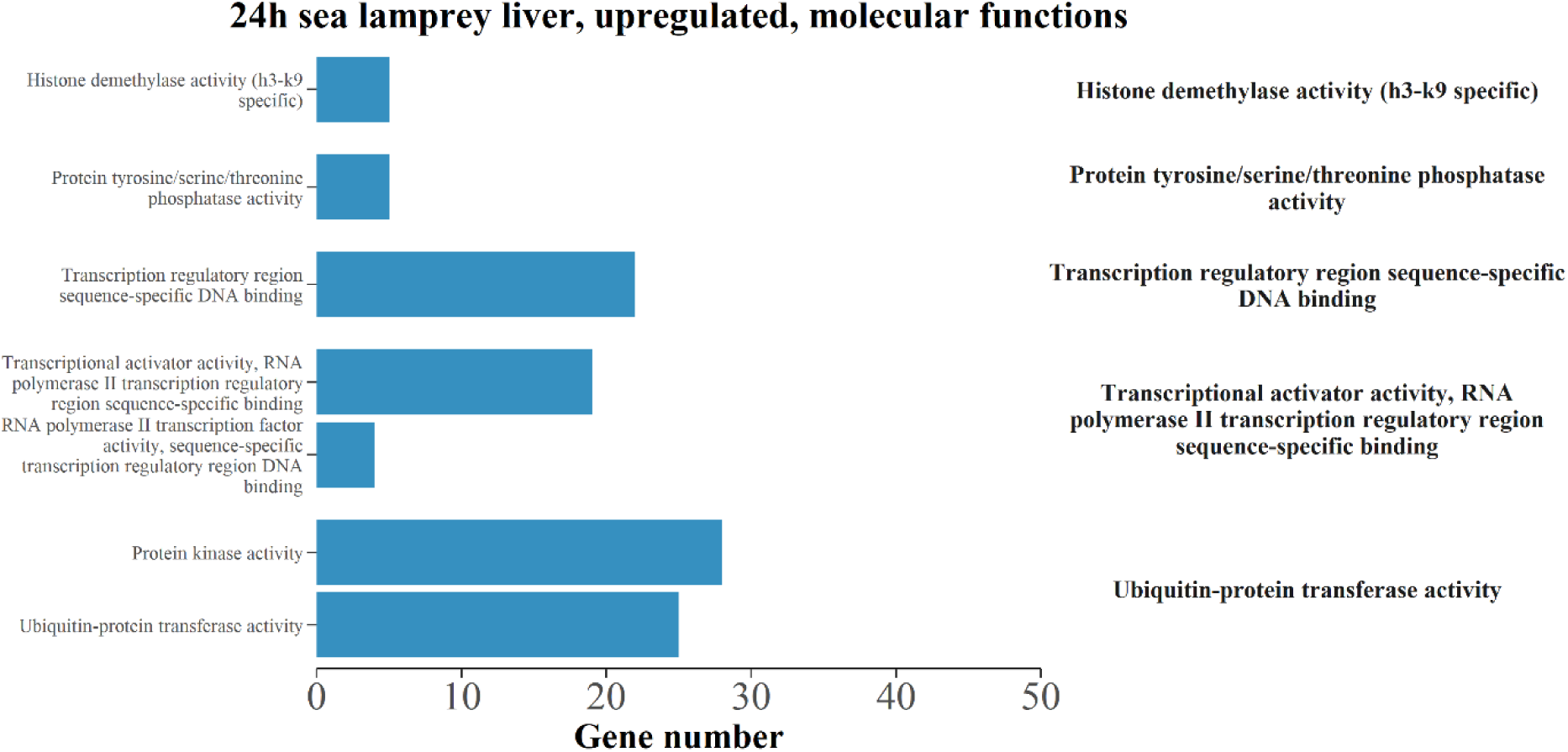
Summary of enriched gene ontology (GO) terms associated with molecular functions in transcripts that were upregulated following 24 h of TFM (3-trifluoromethyl-4’-nitrophenol; 2.21 mg L^-1^ nominal) exposure in the liver of sea lamprey (*Petromyzon marinus*) larvae. Transcripts were considered differentially regulated at a false discovery rate < 0.05. Only GO terms from the functional analysis with an adjusted p < 0.05 and at least four transcripts were considered significantly enriched. REVIGO was used to summarize GO terms to reduce redundancy and group according to similarity (right labels).

**Figure S6:**
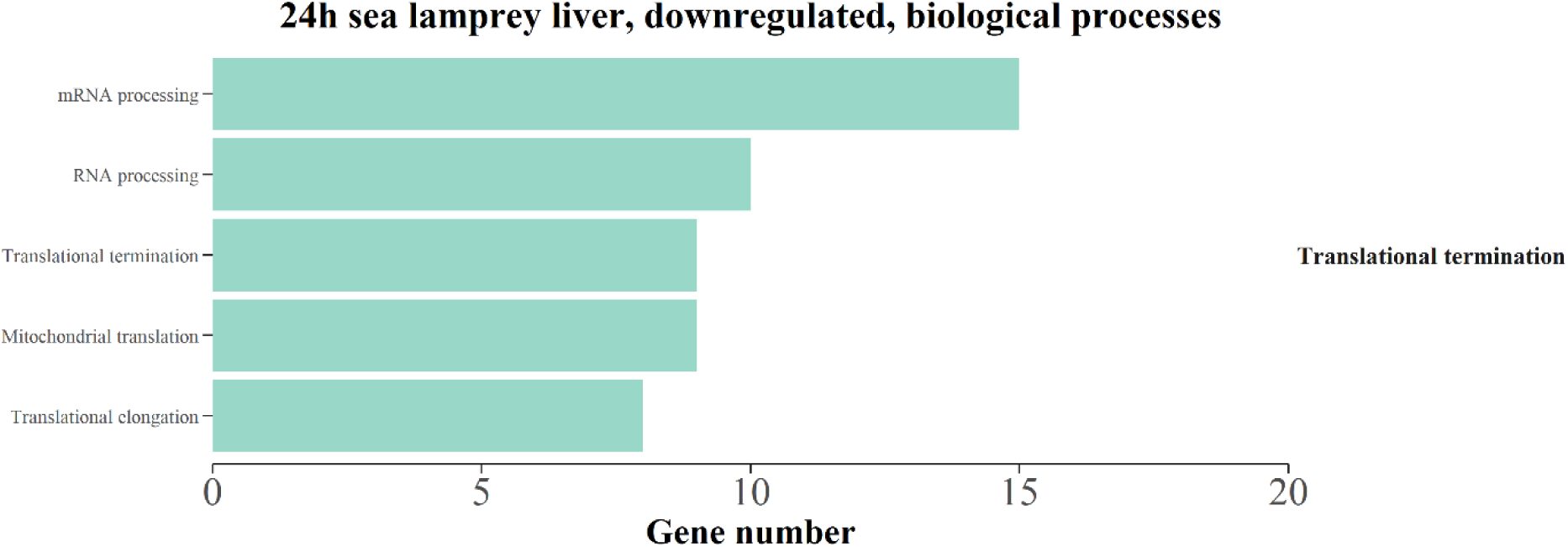
Summary of enriched gene ontology (GO) terms associated with biological processes in transcripts that were downregulated following 24 h of TFM (3-trifluoromethyl-4’-nitrophenol; 2.21 mg L^-1^ nominal) exposure in the liver of sea lamprey (*Petromyzon marinus*) larvae. Transcripts were considered differentially regulated at a false discovery rate < 0.05. Only GO terms from the functional analysis with an adjusted p < 0.05 and at least four transcripts were considered significantly enriched. REVIGO was used to summarize GO terms to reduce redundancy and group according to similarity (right labels).

**Figure S7:**
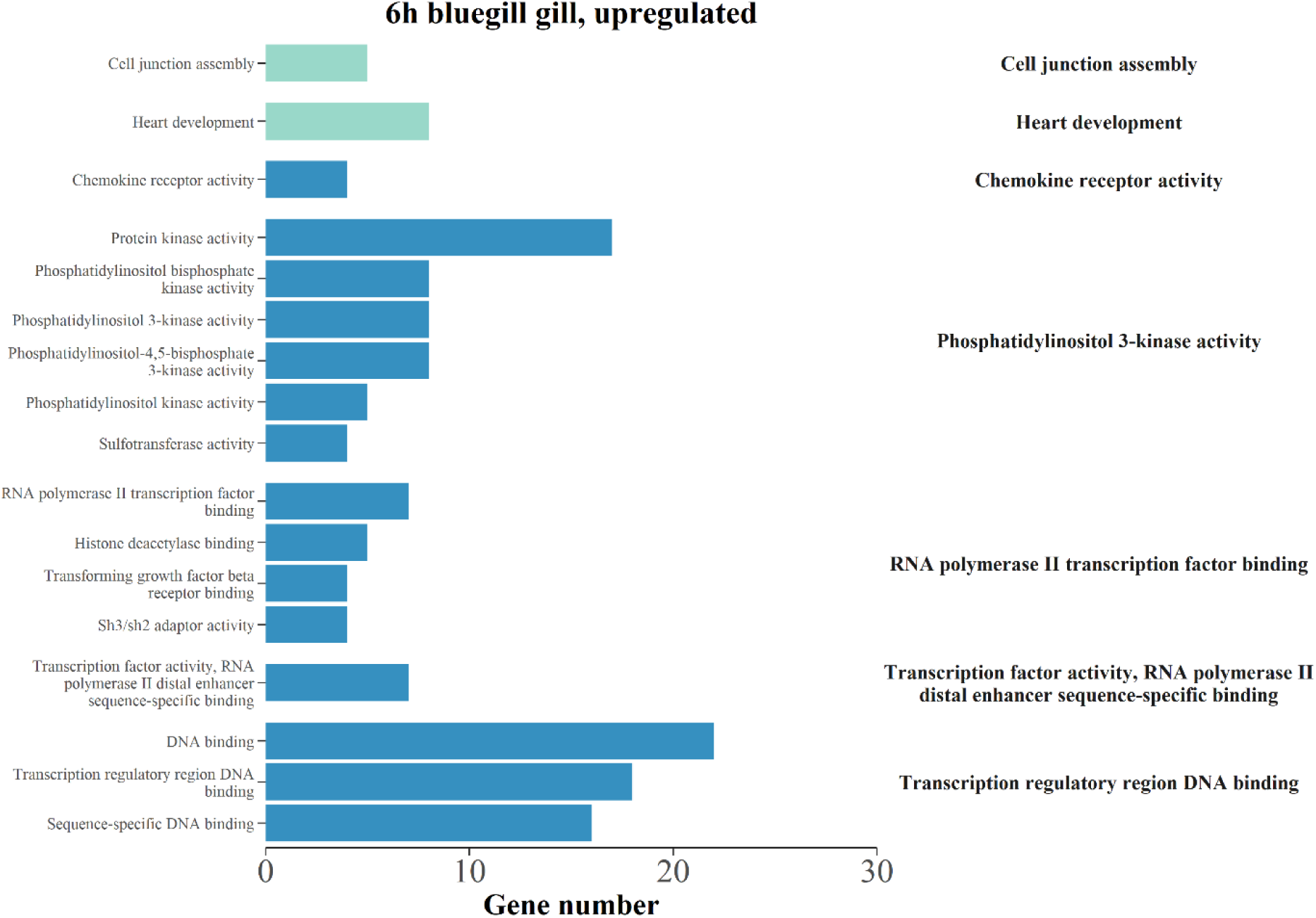
Summary of enriched gene ontology (GO) terms associated with biological processes (green) and molecular functions (blue) in transcripts that were upregulated following 6 h of TFM (3-trifluoromethyl-4’-nitrophenol; 22.06 mg L^-1^ nominal) exposure in the gills of bluegill (*Lepomis macrochirus*). Transcripts were considered differentially regulated at a false discovery rate < 0.05. Only GO terms from the functional analysis with an adjusted p < 0.05 and at least four transcripts were considered significantly enriched. REVIGO was used to summarize GO terms to reduce redundancy and group according to similarity (right labels).

**Figure S8:**
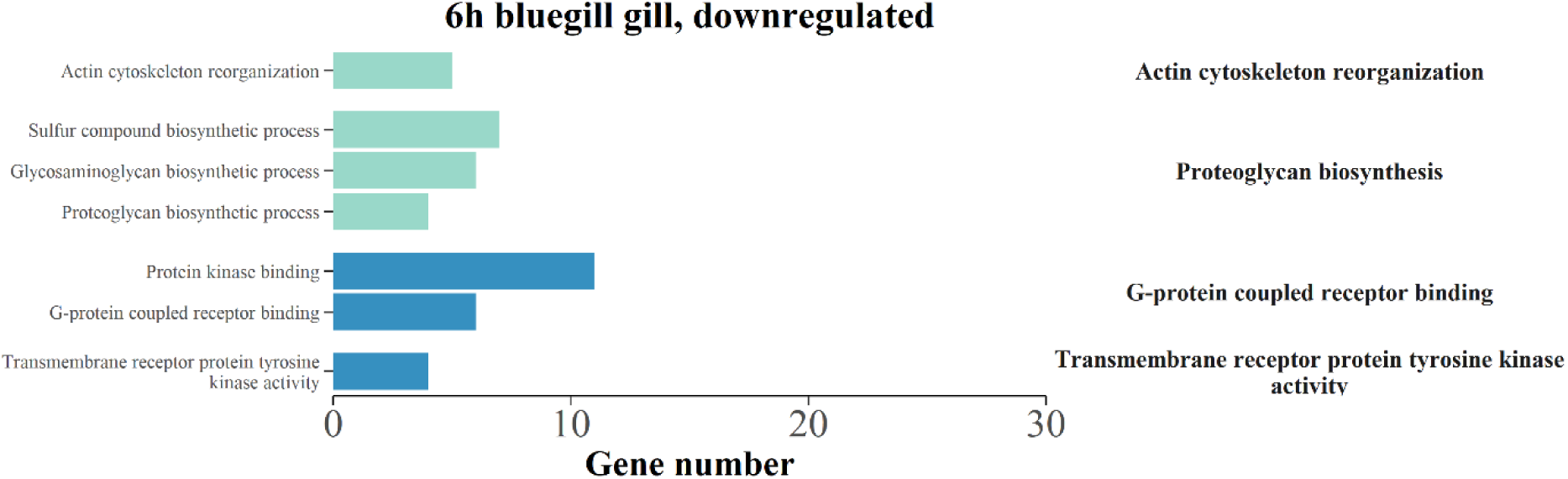
Summary of enriched gene ontology (GO) terms associated with biological processes (green) and molecular functions (blue) in transcripts that were downregulated following 6 h of TFM (3-trifluoromethyl-4’-nitrophenol; 22.06 mg L^-1^ nominal) exposure in the gills of bluegill (*Lepomis macrochirus*). Transcripts were considered differentially regulated at a false discovery rate < 0.05. Only GO terms from the functional analysis with an adjusted p < 0.05 and at least four transcripts were considered significantly enriched. REVIGO was used to summarize GO terms to reduce redundancy and group according to similarity (right labels).

**Figure S9:**
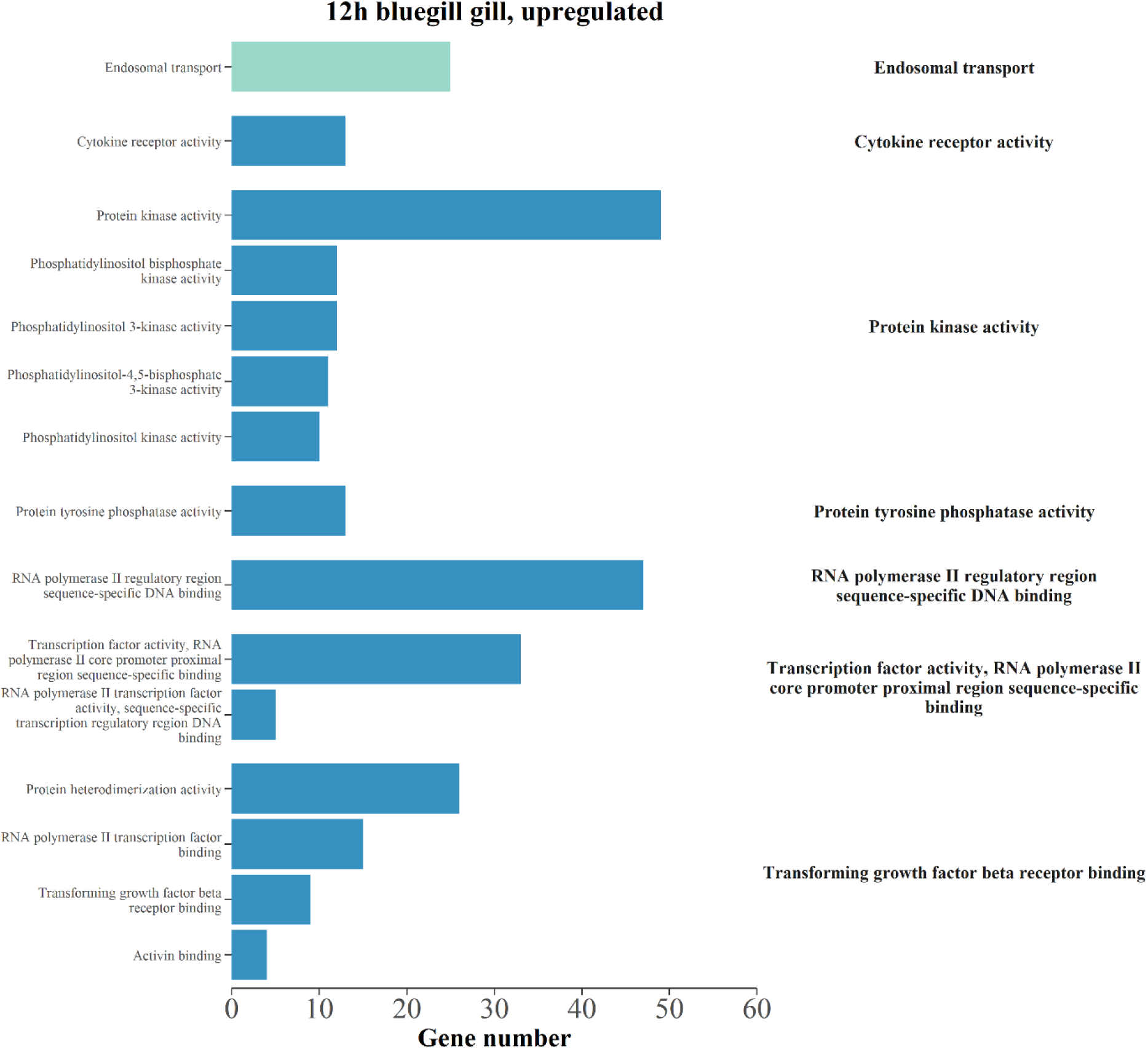
Summary of enriched gene ontology (GO) terms associated with biological processes (green) and molecular functions (blue) in transcripts that were upregulated following 12 h of TFM (3-trifluoromethyl-4’-nitrophenol; 22.06 mg L^-1^ nominal) exposure in the gills of bluegill (*Lepomis macrochirus*). Transcripts were considered differentially regulated at a false discovery rate < 0.05. Only GO terms from the functional analysis with an adjusted p < 0.05 and at least four transcripts were considered significantly enriched. REVIGO was used to summarize GO terms to reduce redundancy and group according to similarity (right labels).

**Figure S10:**
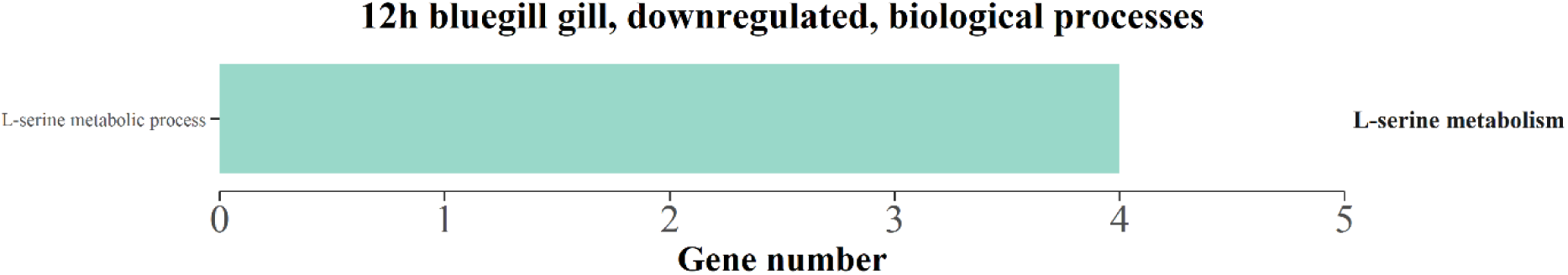
Summary of enriched gene ontology (GO) terms associated with biological processes in transcripts that were downregulated following 12 h of TFM (3-trifluoromethyl-4’- nitrophenol; 22.06 mg L^-1^ nominal) exposure in the gills of bluegill (*Lepomis macrochirus*). Transcripts were considered differentially regulated at a false discovery rate < 0.05. Only GO terms from the functional analysis with an adjusted p < 0.05 and at least four transcripts were considered significantly enriched. REVIGO was used to summarize GO terms to reduce redundancy and group according to similarity (right labels).

**Figure S11:**
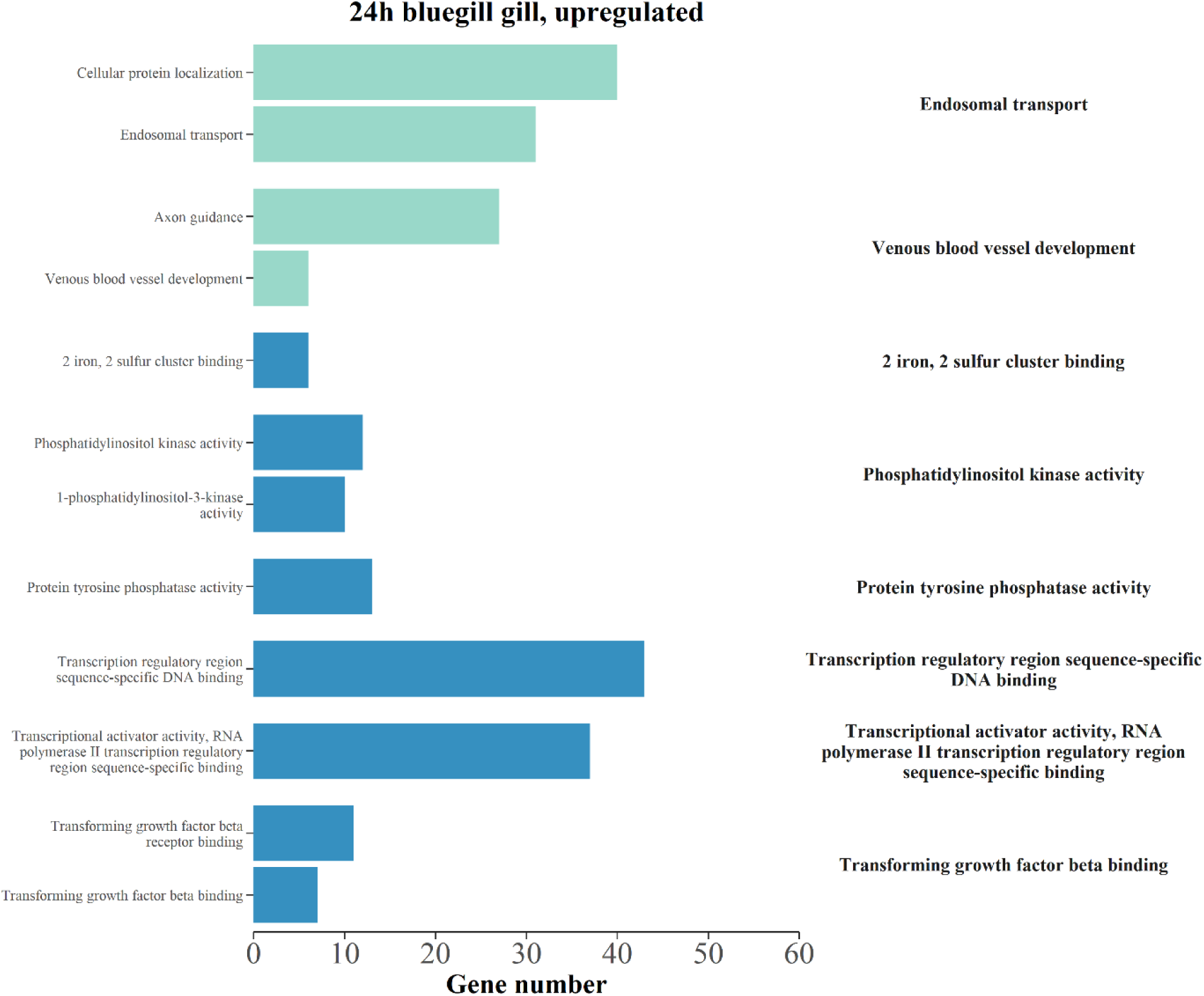
Summary of enriched gene ontology (GO) terms associated with biological processes (green) and molecular functions (blue) in transcripts that were upregulated following 24 h of TFM (3-trifluoromethyl-4’-nitrophenol; 22.06 mg L^-1^ nominal) exposure in the gills of bluegill (*Lepomis macrochirus*). Transcripts were considered differentially regulated at a false discovery rate < 0.05. Only GO terms from the functional analysis with an adjusted p < 0.05 and at least four transcripts were considered significantly enriched. REVIGO was used to summarize GO terms to reduce redundancy and group according to similarity (right labels).

**Figure S12:**
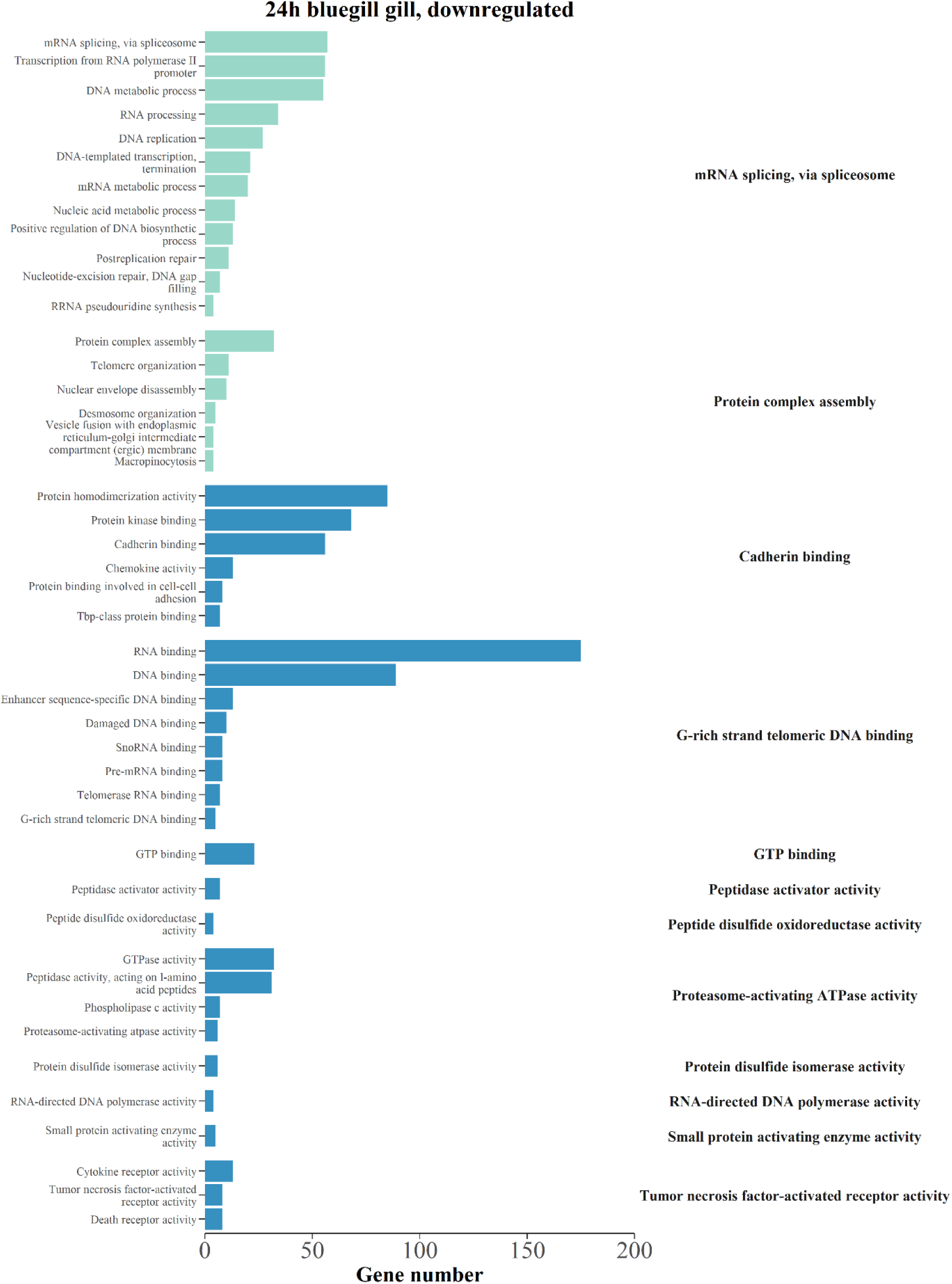
Summary of enriched gene ontology (GO) terms associated with biological processes (green) and molecular functions (blue) in transcripts that were downregulated following 24 h of TFM (3-trifluoromethyl-4’-nitrophenol; 22.06 mg L^-1^ nominal) exposure in the gills of bluegill (*Lepomis macrochirus*). Transcripts were considered differentially regulated at a false discovery rate < 0.05. Only GO terms from the functional analysis with an adjusted p < 0.05 and at least four transcripts were considered significantly enriched. REVIGO was used to summarize GO terms to reduce redundancy and group according to similarity (right labels).

**Figure S13:**
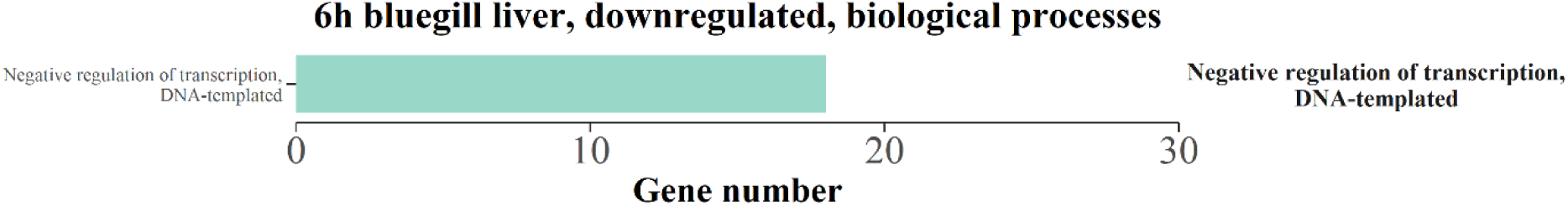
Summary of enriched gene ontology (GO) terms associated with biological processes in transcripts that were downregulated following 6 h of TFM (3-trifluoromethyl-4’- nitrophenol; 22.06 mg L^-1^ nominal) exposure in the liver of bluegill (*Lepomis macrochirus*). Transcripts were considered differentially regulated at a false discovery rate < 0.05. Only GO terms from the functional analysis with an adjusted p < 0.05 and at least four transcripts were considered significantly enriched. REVIGO was used to summarize GO terms to reduce redundancy and group according to similarity (right labels).

**Figure S14:**
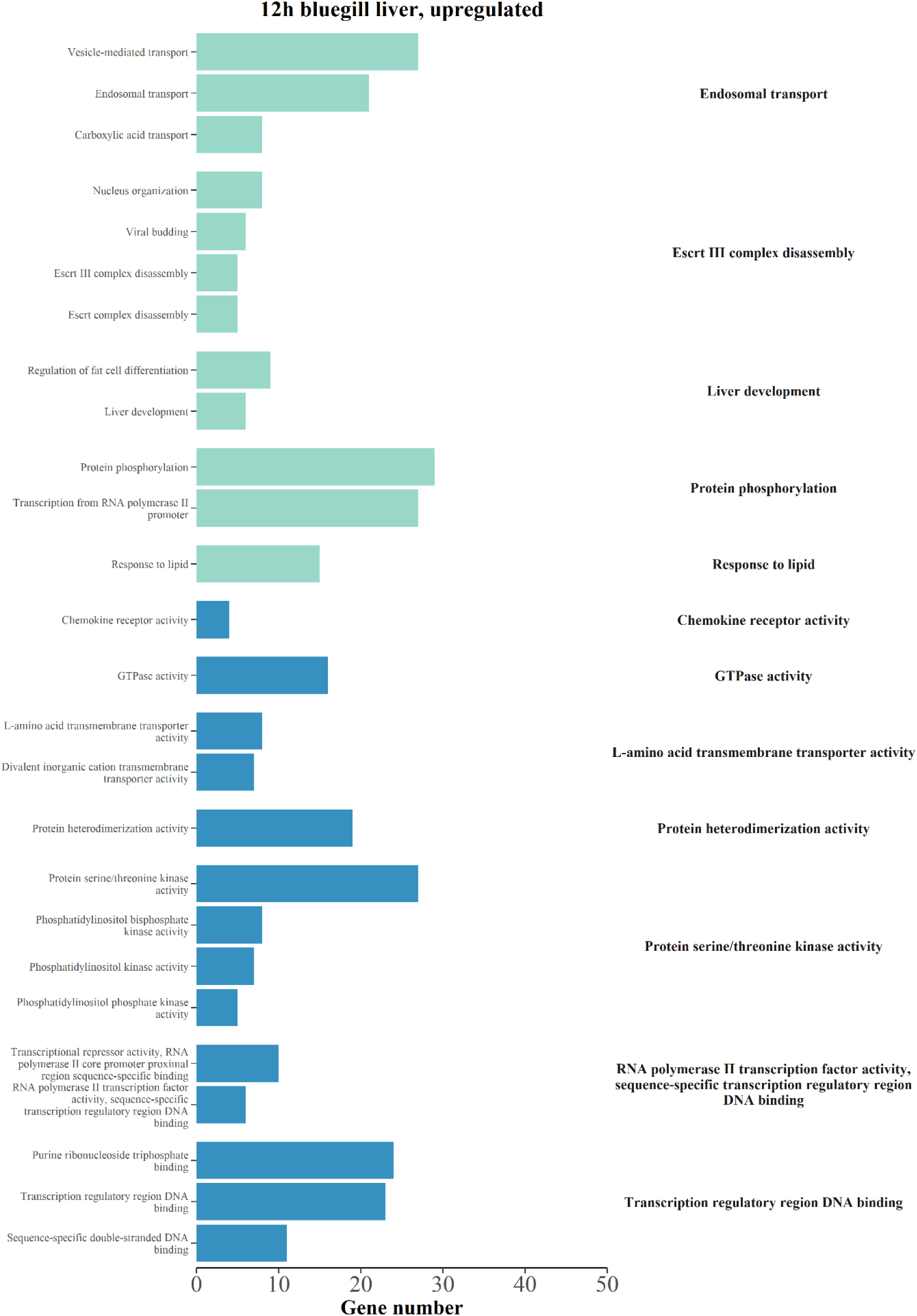
Summary of enriched gene ontology (GO) terms associated with biological processes (green) and molecular functions (blue) in transcripts that were upregulated following 12 h of TFM (3-trifluoromethyl-4’-nitrophenol; 22.06 mg L^-1^ nominal) exposure in the liver of bluegill (*Lepomis macrochirus*). Transcripts were considered differentially regulated at a false discovery rate < 0.05. Only GO terms from the functional analysis with an adjusted p < 0.05 and at least four transcripts were considered significantly enriched. REVIGO was used to summarize GO terms to reduce redundancy and group according to similarity (right labels).

**Figure S15:**
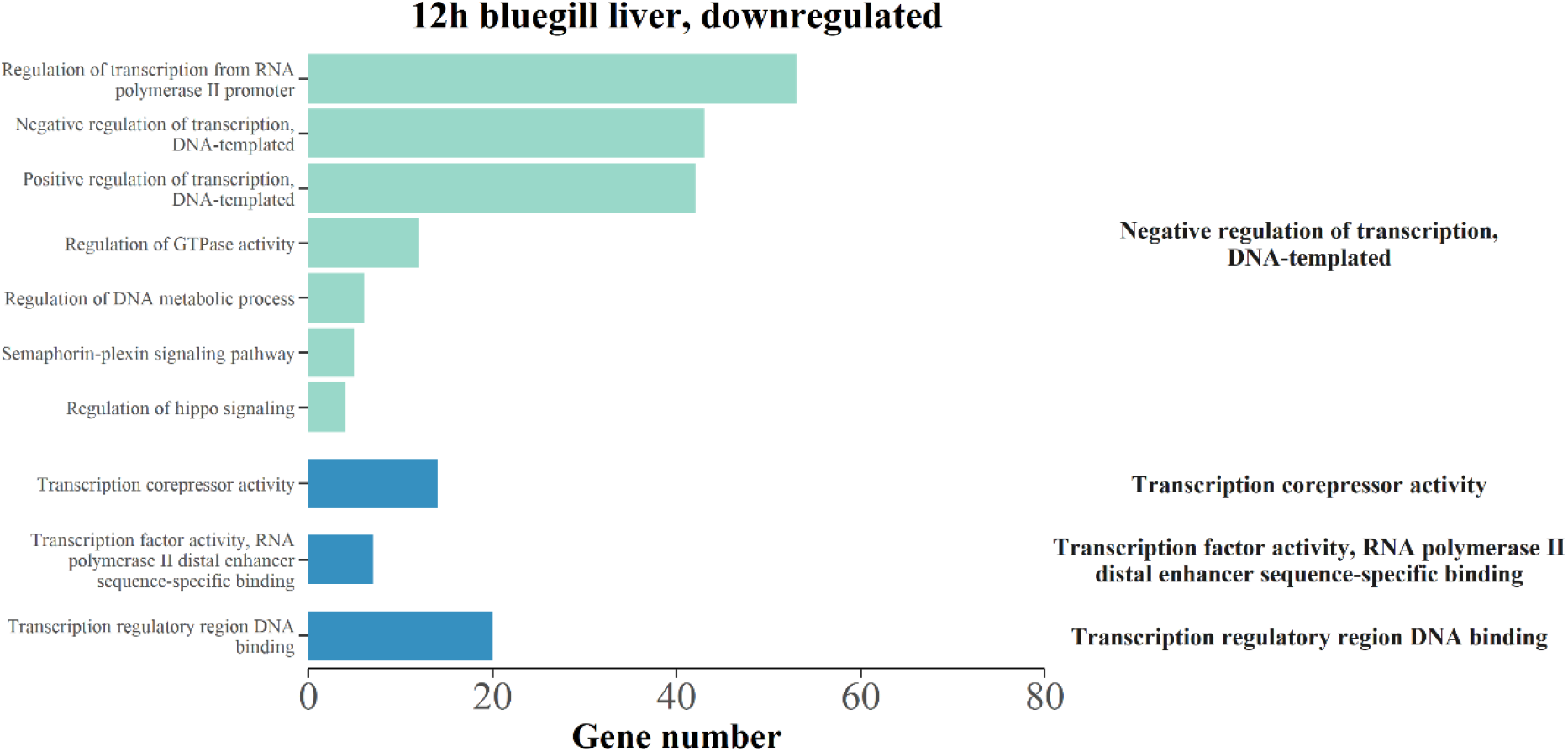
Summary of enriched gene ontology (GO) terms associated with biological processes (green) and molecular functions (blue) in transcripts that were downregulated following 12 h of TFM (3-trifluoromethyl-4’-nitrophenol; 22.06 mg L^-1^ nominal) exposure in the liver of bluegill (*Lepomis macrochirus*). Transcripts were considered differentially regulated at a false discovery rate < 0.05. Only GO terms from the functional analysis with an adjusted p < 0.05 and at least four transcripts were considered significantly enriched. REVIGO was used to summarize GO terms to reduce redundancy and group according to similarity (right labels).

**Figure S16:**
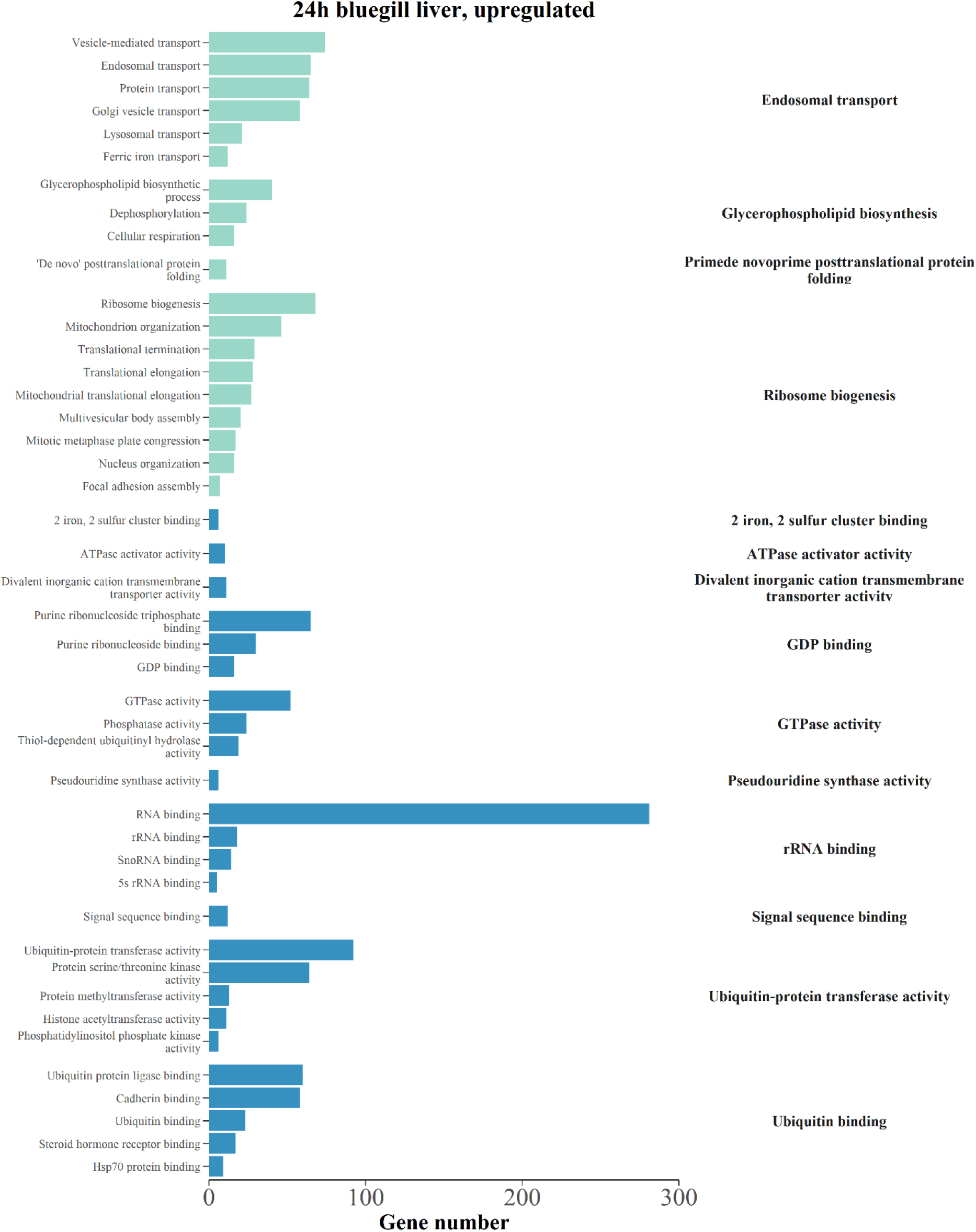
Summary of enriched gene ontology (GO) terms associated with biological processes (green) and molecular functions (blue) in transcripts that were upregulated following 24 h of TFM (3-trifluoromethyl-4’-nitrophenol; 22.06 mg L^-1^ nominal) exposure in the liver of bluegill (*Lepomis macrochirus*). Transcripts were considered differentially regulated at a false discovery rate < 0.05. Only GO terms from the functional analysis with an adjusted p < 0.05 and at least four transcripts were considered significantly enriched. REVIGO was used to summarize GO terms to reduce redundancy and group according to similarity (right labels).

**Figure S17:**
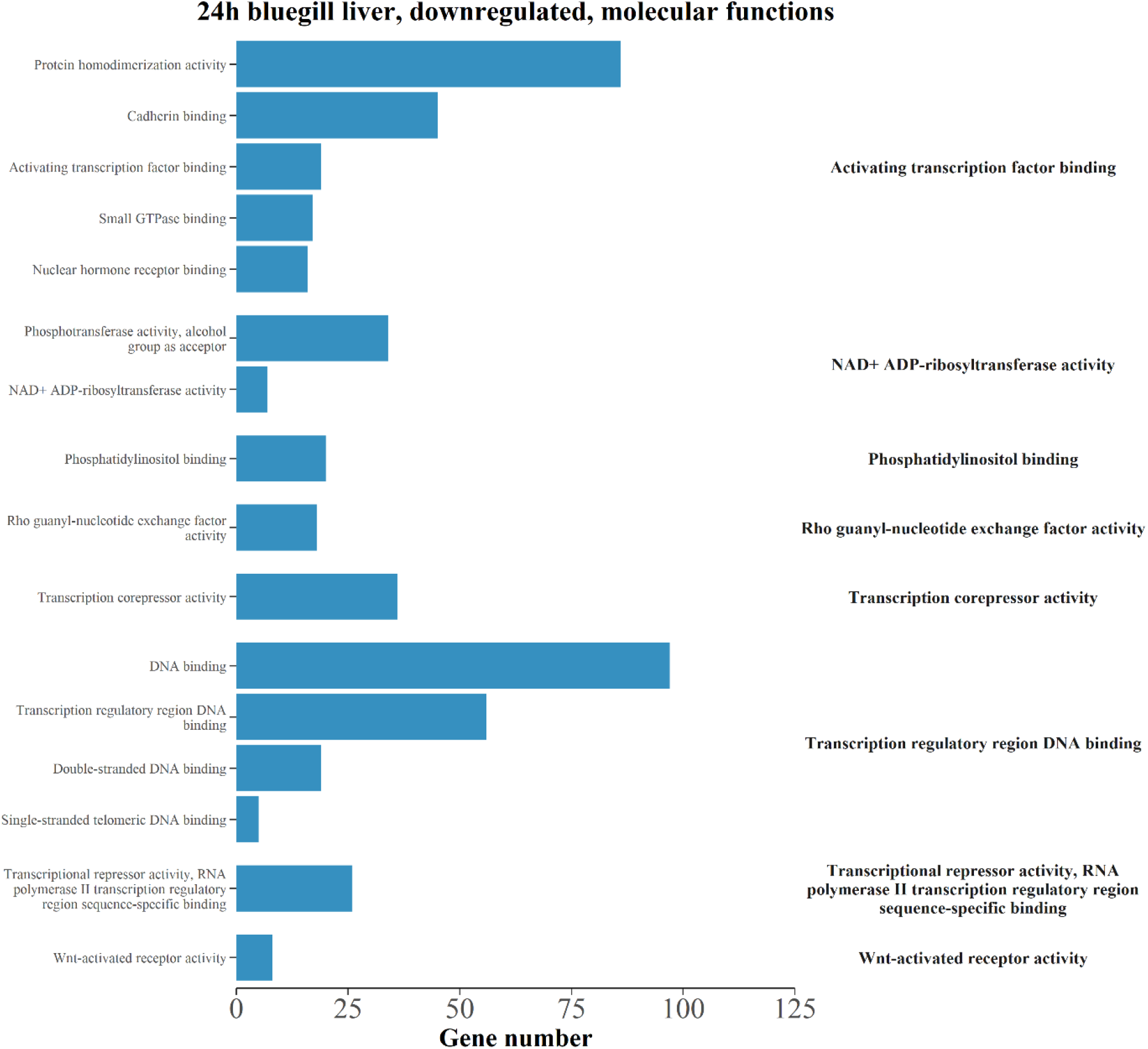
Summary of enriched gene ontology (GO) terms associated with molecular functions in transcripts that were downregulated following 24 h of TFM (3-trifluoromethyl-4’- nitrophenol; 22.06 mg L^-1^ nominal) exposure in the liver of bluegill (*Lepomis macrochirus*). Transcripts were considered differentially regulated at a false discovery rate < 0.05. Only GO terms from the functional analysis with an adjusted p < 0.05 and at least four transcripts were considered significantly enriched. REVIGO was used to summarize GO terms to reduce redundancy and group according to similarity (right labels).

**Table S1:**
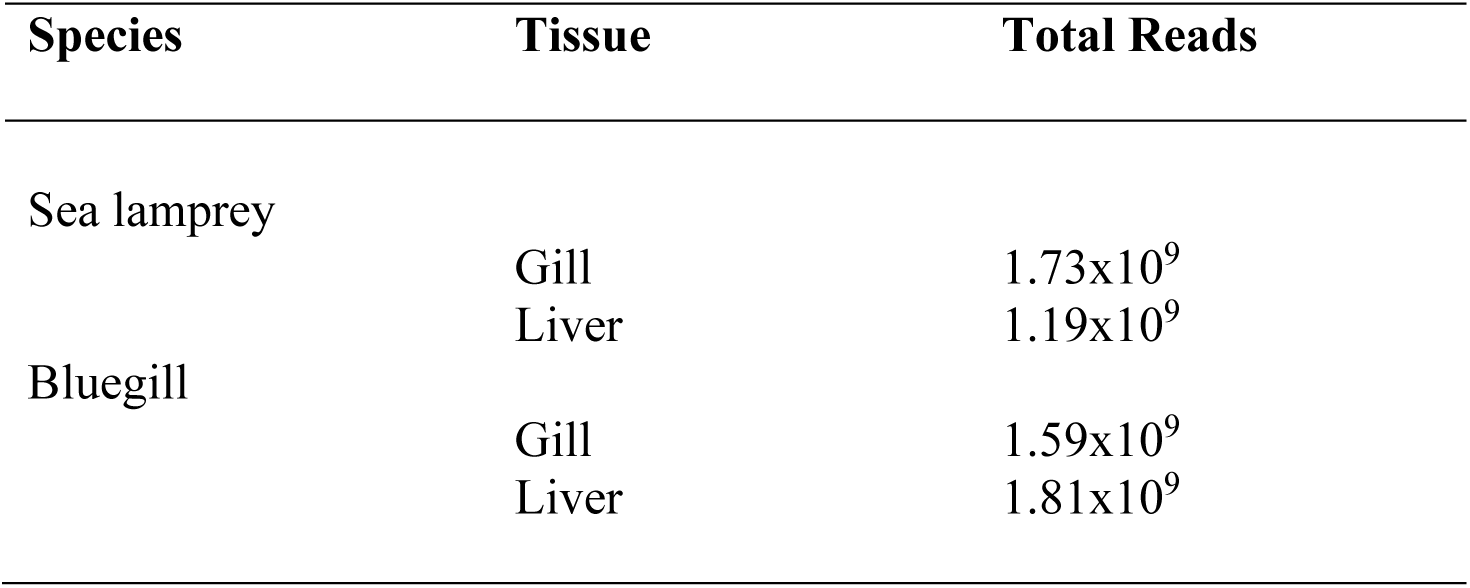
Total sequencing reads counts by species and tissue.

**Table S2.** BUSCO reports comparing the initial Trinity assembly to the Corset-Lace assembly

| Species | Assembly | Complete<br>BUSCOs<br>(C) = S+D | Complete<br>Single-Copy<br>BUSCOs (S) | Complete<br>duplicated<br>BUSCOs (D) | Fragmented<br>BUSCOs (F) | Missing<br>BUSCOs (M) |
| --- | --- | --- | --- | --- | --- | --- |
| *Bluegill* | Trinity | 98.4% | 39.6% | 58.8% | 0.4% | 1.2% |
|  | Corset/Lace | 84.7% | 82.0% | 2.7% | 9.4% | 5.9% |
| *Sea lamprey* | Trinity | 96.1% | 35.7% | 60.4% | 2.4% | 1.5% |
|  | Corset/Lace | 91% | 87.5% | 3.5% | 6.3% | 2.7% |

